# Balanced internal hydration discriminates substrate binding to respiratory complex I

**DOI:** 10.1101/572404

**Authors:** Murilo Hoias Teixeira, Guilherme Menegon Arantes

## Abstract

Molecular recognition of the amphiphilic electron carrier ubiquinone (Q) by respiratory complexes is a fundamental part of electron transfer chains in mitochondria and many bacteria. The primary respiratory complex I binds Q in a long and narrow protein chamber to catalyse its reduction. But, the binding mechanism and the role of chamber hydration in substrate selectivity and stability are unclear. Here, large-scale atomistic molecular dynamics simulations and estimated free energy profiles are used to characterize in detail the binding mechanism to complex I of Q with short and with long isoprenoid tails. A highly stable binding site with two different poses near the chamber exit and a secondary reactive site near the N2 iron-sulfur cluster are found which may lead to an alternative Q redox chemistry and help to explain complex I reactivity. The binding energetics depends mainly on polar interactions of the Q-head and on the counterbalanced hydration of Q-tail isoprenoid units and hydrophobic residues inside the protein chamber. Selectivity upon variation of tail length arises by shifting the hydration balance. This internal hydration mechanism may have implications for binding of amphiphilic molecules to cavities in other membrane proteins.

## Introduction

Respiratory complex I also known as NADH:ubiquinone oxidoreductase is the main entry protein of electron transfer chains in the inner membrane of mitochondria and in many bacteria. It catalyzes oxidation of nicotine adenine dinucleotide (NADH) through a flavin mononucleotide and a chain of iron-sulfur (FeS) clusters to reduce ubiquinone (Q), an amphiphile composed by a *p*-benzoquinone ring (Q-head) attached to an isoprenoid chain (Q-tail), which diffuses along the membrane and carries electrons to subsequent respiratory complexes. Complex I is also a reversible proton pump and couples the redox process with generation of an electrochemical gradient across the membrane, thus contributing to ATP synthesis^1–5^. As an essential metabolic enzyme and a primary site for production of reactive oxygen species, malfunction of complex I has been linked to several common neuromuscular, degenerative and metabolic diseases, ischemia-reperfusion injury and aging^6,7^.

Complex I is one of the largest asymmetrical membrane proteins known. The prokaryotic enzyme is usually composed by 14 core subunits (550 kDa mass) containing all the redox centers and proposed proton pumping channels. These subunits are sufficient for catalysis and highly conserved from bacteria to human enzymes. The mammalian complex I has 31 additional supernumerary subunits (total of 45 subunits and 980 kDa mass) involved in complex assembly, stability and specialized metabolic roles.

The first entire atomic structure of complex I was determined^8^ for the eubacterium *Thermus thermophilus* enzyme by X-ray crystallography with a resolution of 3.3Å. The L-shaped structure is composed by a membrane-bound arm where the proton channels are located, and a hydrophilic peripheral arm where all the redox cofactors and NÅDH binding site are found (Figure 1A). Mitochondrial structures from the yeast *Yarrowia lipolytica*^9^ and from several mammals (bovine,^10^ ovine,^11^ porcine^12^ and mouse^13^) determined more recently by cryo-EM experiments revealed the external location of supernumerary subunits forming a protective shell around the core. The overall geometry of the core subunits is similar in all determined structures which suggests conservation of Q binding and catalytic mechanisms among species.

**Figure 1:**
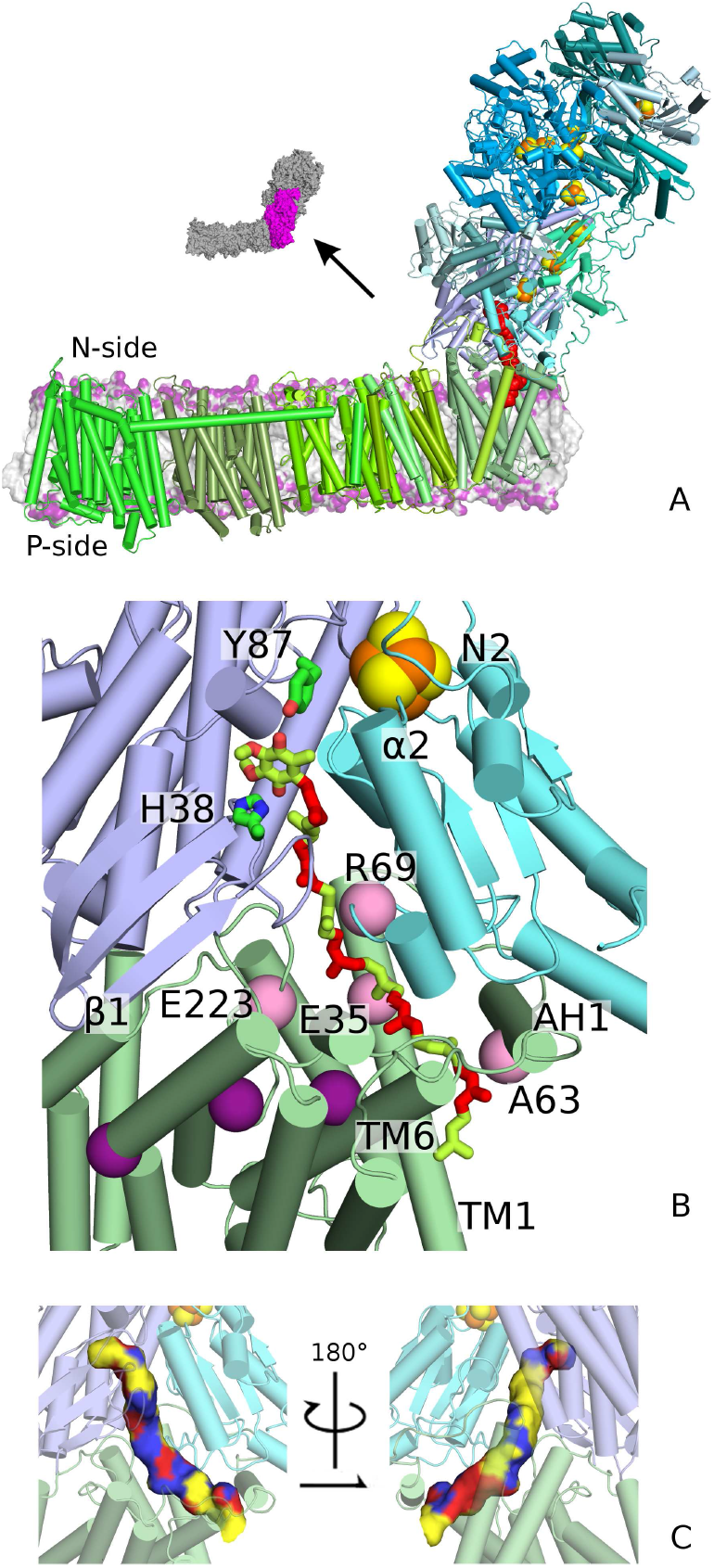
Structure of the respiratory complex I and its Q binding chamber. (A) Overview in cartoon of the structural model from *T. thermophilus* embedded in a lipid membrane. FeS clusters are shown as orange and yellow spheres. Q_10_ is modeled in the reactive site in red. Inset shows the position of subunits Nqo4, Nqo6 and Nqo8. (B) Close view of subunits Nqo4 in light blue, Nqo6 in cyan and Nqo8 in pale green that form the Q-chamber with Q_10_ colored in green and red alternating for each isoprenoid unit. Elements of secondary structure and residues (green sticks and pink spheres) discussed in the text are indicated. Purple spheres show Nqo8 residues Glu163, Glu213 and Glu248 of the proposed E-channel.^8^ (C) Molecular surface of bound Q colored by the first solvation layer: yellow is a hydrophobic residue, red is a hydrophilic residue and blue is water.

A 35 Å long and narrow Q binding chamber was identified on the interface of subunits Nqo4 (49kDa in bovine nomenclature), Nqo6 (PSST) and Nqo8 (ND1) in the *T. thermophilus* structure (Fig. 1B). This remarkable protein cavity (Q-chamber) starts on top of a cleft between subunits Nqo4 and Nqo6 where the substrate Q-head may bind, receive electrons from the nearby N2 FeS cluster (center-to-center distance ~13 Å) and form hydrogen bonds through its carbonyl oxygens to the side chains of Nqo4 Tyr87 and His38, which are both invariant and required for full enzymatic activity. X-ray data show the inhibitor piericidin A and the substrate analogue decylubiquinone, both containing short hydrophobic tails, bind in the Q-chamber at this position.^8,9^ The chamber continues through an amphipathic region, roughly composed by a charged surface facing the membrane arm and by a large hydrophobic patch on the opposite side (Figs. 1B and C). It ends exposed to the lipid membrane in an exit formed by Nqo8 helices TM1, TM6 and amphipathic helix AH1. The Q-chamber geometry and the free volume available to accommodate Q inside are similar among all determined active structures.^8–11,13^

Although complex I has been cocrystallized with small Q analogs,^8,9^ it is still unclear how natural substrates with long isoprenoid chains such as ubiquinone-10 (Q_10_) will bind and how many stable binding sites or poses may be present inside the long Q-chamber. Will each putative binding site lead to different Q redox chemistry? Is binding dominated by Q-head or by Q-tail interactions? Does water penetrate in the Q-chamber and play a role during binding? Answers will help to understand the mechanism of substrate binding in the amphipathic Q-chamber and to design more potent and selective inhibitors.

Molecular simulation is a valuable tool to investigate binding pathways and energetics of ligands complexed to protein tunnels or cavities.^14–19^ For instance, unbinding of a small ligand from the prototypical T4 lysozyme engineered cavity has been shown to proceed through multiple competitive tunnels in the protein.^19^ Pathways of plastoquinone exchange along different channels in the photosystem II complex have been determined by coarse-grain models^20^. For the respiratory complex I, a recent simulation study^21^ explored how the Q redox state modulates binding in the Q-chamber and triggers the coupled proton pumping.

Here we present large scale molecular dynamics simulations and estimate free energy profiles for binding of Q with long (Q_10_) and short (ubiquinone-2, Q_2_) isoprenoid chains to the hydrated chamber inside complex I from *T. thermophilus.* Notably, our model is based on the entire complex I structure^8^ with corrected coordinates for Nqo6 loop *α2–β*2 and on a calibrated force field for Q,^22^ as described in the next section. Results show Q is stable in two separated binding and reactive sites, respectively near the Q-chamber exit and cluster N2. The binding site (BS) is broad with at least two different stable poses. The binding mechanism depends on an interesting but previously uncharacterized interplay of Q-head interactions and hydration of both Q-tail and internal chamber residues. We discuss the implications for substrate selectivity and reactivity catalysed by complex I and conlcude this binding mechanism may be employed by amphiphilic molecules binding to other membrane proteins.

## Computational Methods

### Set-up of protein model and molecular dynamics simulations

The X-ray structure of the complete respiratory complex I from *T. thermophilus* (PDB ID: 4HEA)^8^ was used to build the protein model. All core subunits (Nqo1-14) and two accessory subunits Nqo15-16 found in species related to *T. Thermophilus* were included. Coordinates for internal segments from subunit Nqo3 (residues 56-72 and 144-147) missing from the PDB file were constructed *de novo* using MODELLER (version 9.15)^23^ with default settings. The Nqo3 residues are distant from the Q-chamber by more than 50A. Nqo6 residues 65-69, also missing from the PDB file, are located in the Q-chamber and in direct contact with bound Q. These residues are part of loop α2–β2 found in a different conformation in the corresponding PSST domain of the resolved active mammalian structures^10,11,13^ (Fig. S2). Thus, we decided to rebuilt the entire loop (segment between residues D55-P72 of Nqo6) with MODELLER but using the equivalent segment (A82-P98 of subunit PSST) from the ovine enzyme (PDB ID: 5LNK)^11^ as a structural template. The other two flexible loops in the Q-chamber (Nqo4 β 1-β2 and Nqo8 TM5–TM6) should be in appropriate position for Q binding as the *T. thermophilus* structure was co-crystalized with a substrate analogue. The rebuilt coordinates are in excellent agreement with a refined model for the *T. Thermophilus* structure (Leonid Sazanov – IST Austria, personal communication) in which coordinates for Nqo6 loop α2–β2, including residues 65-69, were rebuilt directly from re-refinement of the original electron density.^8^

Protonation states of side-chains were adjusted to neutral pH (positive charge for K and R, negative for D and E, and neutral for all other residues), except for Nqo4 His38 which was protonated. All FeS centers were modeled in their reduced form. Equivalent charge states were used in previous simulations.^24^

Ubiquinone was modeled in the oxidized form with 10 (Q_10_) or 2 (Q_2_) isoprenoid units. Q_2_ was first docked in the reactive position near Nqo4 Tyr87 and His38 using AutoDock 4.^25^ The remaining isoprenoid units for Q_10_ were manually placed along the Q-chamber^8^ in an extended conformation. The last two isoprenoid units in Q_10_ pass through the least narrow Q-chamber exit formed between Nqo8 helices TM1, TM6 and AH1 and are exposed to the lipid membrane. All protein atoms remained in their X-ray structure as the Q-chamber has enough free volume to accommodate Q without protein movements.

The protein complex was embedded^26^ in a solvated POPC (1-palmitoyl-2-oleoyl-sn-glycero-3-phosphocholine) membrane with 1036 lipid molecules, 192951 water molecules, and 650 Na+ and 607 Cl^-^ ions to neutralize the total system charge and keep a 0.1 M salt concentration, resulting in a total of 794102 atoms. This model was relaxed during two molecular dynamics simulations of 100 ns each, first with all protein heavy atoms tethered to their initial position by harmonic restraints, then with all atoms free to move. This procedure resulted in the initial structural model used for the free energy simulations.

Modelling of the Q binding process requires a balanced description of Q interactions with the hydrated protein interior and the lipid membrane. This was accomplished by using a calibrated force-field for Q, previously shown to give very good agreement between simulated and experimental water-to-membrane partition coefficients and free energies.^22^ Interactions of protein, lipids and ions were described with the all-atom CHARMM36 force-field.^27,28^ Water was represented by standard TIP3P.^29^ FeS centers were described using the Chang & Kim^30^ parameters with corrections proposed by McCullagh & Voth.^31^ These corrections are important to keep the cuboidal structure in Fe_4_S_4_ clusters.

All molecular dynamics simulations were performed with GROMACS (versions 5.1.3 and 20 1 6.3)^32^ at constant temperature of 310 K, pressure of 1 atm and a time step of 2 fs. Long-range electrostatics was treated with the Particle Mesh Ewald method.^33^ Further details are given in the Electronic Supplementary Information (ESI). Visualization and figure plotting were done using PyMol^34^ and Matplotlib.^35^ All simulation data and workflow scripts are available from the authors upon request.

### Reaction coordinates and free-energy calculations

Two reaction coordinates were used to describe the Q binding process: the distance between the centers of mass (dCOM) of the six carbon atoms in the Q ring and ten C_α_ of residues in subunit Nqo4 surrounding the Q-head in the reactive site (Fig. S1); and a pathway collective variable (Path CV)^36^ using a distance metric^16,17^ as implemented in PLUMED 2.3.1^37^ between the Q-head and 55 C_α_ of residues in subunits Nqo4, Nqo6 and Nqo8 (Fig. S1) in direct contact with Q inside the chamber. The Path CV was evaluated with respect to 30 milestone configurations representing progressive binding of Q along the chamber (see ESI Methods). These two kinds of reaction coordinates have already been used successfully to describe ligand binding along protein tunnels or cavities.^16–19^ Results are presented using the shifted coordinate sdCOM = 4 nm — dCOM, so that entrance of Q from the membrane into the Q-chamber proceeds from low to high values and is seen from left to right in the figures shown. This representation is more intuitive than using dCOM or Path CV, which run from high to low values during entrance of Q. All the conclusions drawn here are equivalent when projections over either dCOM (and its sdCOM representation) or the Path CV are used to describe the results (Fig. S5 and ESI).

Free energy profiles for Q binding were estimated with umbrella sampling (US) simulations.^38^ Initial configurations for each umbrella window were generated from the initial structural models described above by pulling the terminal carbon in the isoprenoid tail along the membrane plane (XY direction) with a pulling velocity of 0.2 m/s. About 50 ns of steered molecular dynamics were enough to drive Q from the initially prepared reactive configuration (dCOM=0.4 nm) towards protein dissociation to the membrane (dCOM=4.1 nm). US windows separated by 0.1 nm were chosen to cover the full range of dCOM, which was restrained with a harmonic potential with force constant *k_umb,dCOM_* = 2000 kJ mol^−1^ nm^−2^. Additional windows were introduced to increase sampling and overlap of the dCOM and Path CV distributions, which were then restrained with force constants *k_umb,dCOM_* = 200 and *k_umb,Path_* = 200 kJ mol^−1^ nm^−2^. A total of 55 and 46 windows were used to emulate binding for Q_10_ and Q_2_, respectively (Table S1). Each coordinate window was sampled for 200 ns for Q_10_ and 150 ns for Q_2_, with a sample collected every 20ps. Total aggregate simulation time was over 28 μs, including preliminary runs shown in the ESI. Potentials of mean-force were obtained from the 2D reaction coordinate distribution with WHAM^38,39^ and the statistical uncertainty was estimated as 95% confidence intervals by bootstrap analysis with 50 resampling steps.^40^ The initial 50 ns of each window were discarded to allow equilibration of orthogonal degrees of freedom. The 1D profiles shown are projections over the minimum 2D free energy pathway. All calculated properties such as number of contacts, hydration, COM distances, etc, were collected over the aggregated windows for each US simulation, binned along the reaction coordinate and averaged for each bin.

## Results

### Q transits through a hydrated chamber with a highly stable binding site

Q binding is described here by the reaction coordinate sdCOM (see Methods and Fig. S1). Entrance of Q from the lipid membrane into complex I corresponds to sdCOM running from low to high values.

Figure 2 shows the free energy profile for Q_10_ binding estimated from umbrella sampling with molecular dynamics simulations. Before entrance of Q, the chamber interior in complex I is filled with tens of water molecules (Fig. S4). To initiate the binding process, a Q_10_ molecule approaching from the membrane pool will exchange contacts with lipids for exposed hydrophobic residues in Nqo8 helices TM1, TM6 and AH1 (denominated CE position, inset I of Fig. 2A. See also Figs. 2E and S3B). The hydrophobic and flexible isoprenoid Q-tail often folds on itself (Fig. S6A) and the average Q-head distance to the membrane center is only 0.2 nm higher than the equilibrium distance when Q is free in the lipid (Fig. 2B). This should minimize the free energy cost associated with formation of the initial protein-Q encounter complex.^22^ The Q-head passes through the hydrophobic chamber exit at sdCOM=0.4-0.6 nm (Fig. S6A) with almost no energy cost as there is enough area for Q transit without significant protein deformation (Fig. S6B).

**Figure 2:**
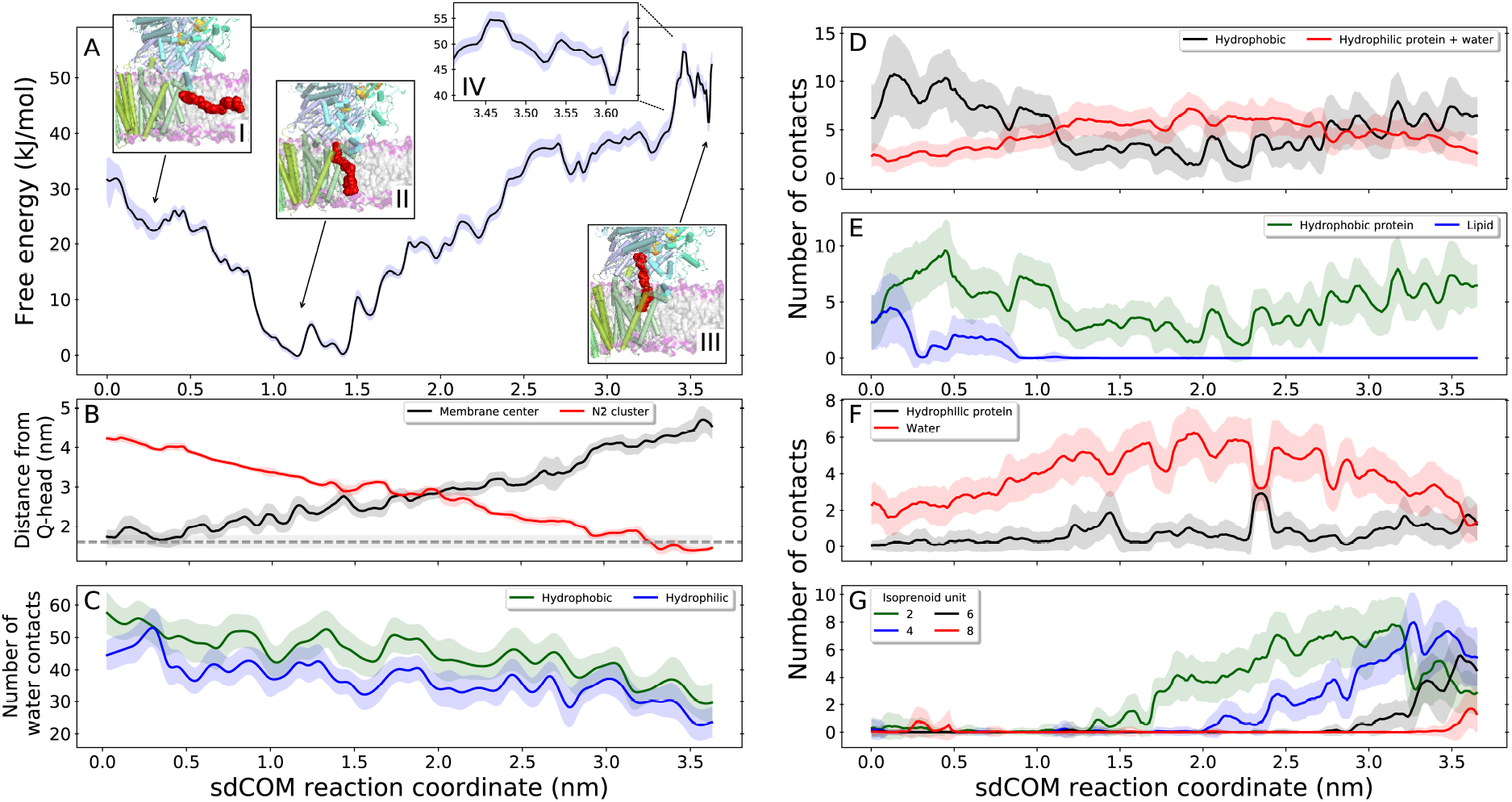
Binding of Q_10_ along the chamber in respiratory complex I. (A) Free energy profile (black line) with statistical uncertainty (blue shadow). Insets I, II and III show the structure of complex I with Q_10_ (red) located in the chamber exit (CE), binding site (BS) and reactive site (RS), respectively. Inset IV zooms profile in the RS region. (B) Distance of Q-head COM to the membrane center (black) and to the N2 cluster COM (red). Dashed line (gray) indicates the Q-head to membrane equilibrium distance when Q is free in the lipid,^22^ 1.60 ± 0.15 nm. (C) Number of water contacts to hydrophobic (green) and hydrophilic (blue) protein residues in the Q-chamber interior. Q oxygen contacts with: (D) hydrophobic (black, apolar protein + lipid hydrocarbon tail) and hydrophilic (red, polar and charged protein + water) groups; (E) hydrophobic protein (green) and lipid (blue, both polar head and hydrophobic tail) groups; (F) hydrophilic protein (black) and water (red) groups. (G) Water contacts with Q_10_ isoprenoid units 2 (green), 4 (blue), 6 (black) and 8 (red). The x-axis in all panels displays the sdCOM reaction coordinate. In panels B-G, lines indicate average properties and shadows indicate one standard deviation.

A broad and highly stable binding site is found between 1.0<sdCOM<1.5 nm (BS position, inset II in Fig. 2A) with two iso-energetic binding poses: at sdCOM=1.1 nm, the Q-head is highly hydrated and at sdCOM=1.4 nm, Q oxygens form hydrogen bonds with conserved Nqo8 Arg36 and Lys65 (Fig. 3A). This is the global minimum of the free energy profile, suggesting that complex I will be frequently loaded with a Q_10_ molecule at this site. Hydrophobic residues internal to the Q-chamber dehydrate due to entrance of Q and partial expulsion of water molecules from the chamber to the aqueous phase (Fig. 2C). Q-head interactions with hydrophobic groups decrease, hydrophilic contacts are formed and hydration increases from an average of 2 water contacts in sdCOM=0.3 nm to almost 5 contacts in sdCOM=1.2 nm (Figs. 2D and 2F).

**Figure 3:**
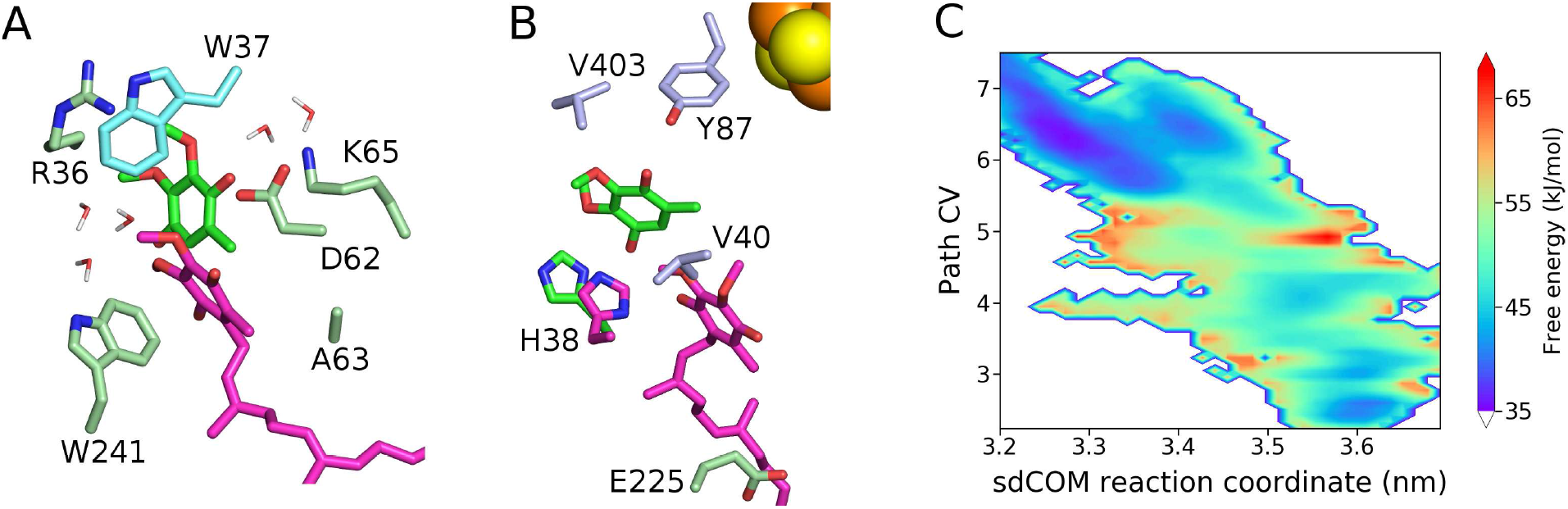
Binding (BS) and reactive (RS) sites in complex I. (A) Protein environment in the BS with two stable Q_10_ positions in magenta (sdCOM=1.1 nm) and green (sdCOM=1.4 nm). Isoprenoid tail of the green Q was removed to ease visualization and water molecules indicate chamber hydration. (B) Environment of Q_10_ at the pre-RS position (sdCOM=3.3 nm, Path CV=6) in magenta and at the lowest minimum in the RS (sdCOM=3.6 nm, Path CV=2.5) in green, with the two respective Nqo4 His38 conformations shown in the same colors. 0ther residues are colored as their subunits in Fig. 1. (C) Two-dimensional free energy profile for Q_10_ binding in the RS region.

As Q_10_ proceeds into the chamber, the free energy increases through a sequence of shallow local minima and reaches a plateau of ~37 kJ/mol between 2.6<sdCOM<3.4 nm (Fig. 2A). Considerable wetting of isoprenoid units 1 to 4 is observed in the same sdCOM range (Fig. 2G).

In the minimum found at 3.2<sdCOM<3.4 nm, a Q carbonyl oxygen can form a hydrogen bond with the protonated Nqo4 His38 side-chain (pre-RS position, Fig. 3B). Q is still far from Nqo4 Tyr87 and the other closest acidic residue, Nqo8 Glu225, is more than 8Å away. In order to form hydrogen bonds with both His38 and Tyr87, the barrier at sdCOM=3.45 nm has to be transposed.

A metastable reactive site (RS position, inset III of Fig. 2A) is found at 3.5<sdCOM<3.65 nm where the lowest free energy is 42 kJ/mol (inset IV of Fig. 2A). The chamber is more hydrophobic in this region (Fig. 1C) and the Q-head dehydrates to form hydrogen bonds with both Tyr87 and His38 (Figs. 2D and 2F). The Q-head is 13Å from the N2 cluster, in appropriate position to be reduced to the quinol form (Fig. 3B). Two other minima may be identified in the RS (using the Path CV reaction coordinate, Fig. 3C), with the Q-head performing only one hydrogen bond with either Tyr87 (Path CV=4) or His38 (Path CV=3), as found in a previous simulation.^41^

There is a network of ionic side chain contacts near the Q-chamber (Fig. S3A). While part of these contacts remain stable during the full binding process (for instance, Nqo8 Arg294 and Lys65. See Fig. S7 and the associated Supplementary Results), some ionic contacts are perturbed either by hydrogen bonding with the Q-head (Nqo8 Arg36 and Nqo6 Arg69) or indirectly (Nqo8 Arg216) via a combination of electrostatic interactions. In the proposed E-channel (Fig. 1B), Nqo8 Glu213 approaches Glu163 at 2.2<sdCOM<3.5 nm in response to coordination with Arg216, which also depends on contacts with Nqo8 Glu248 and the highly flexible Glu223 (Fig. S7). These correlated motions might link Q transit with activation of the E-channel and proton pumping in the membrane arm.

### A shorter isoprenoid tail increases stability of Q in the reactive site

Although natural complex I substrates have 6 to 10 isoprenoid units, experiments are often carried out with more soluble short-tail analogues.^1,42–45^ The effect of Q-tail length on binding was investigated by simulating Q_2_ as shown in Fig. 4. The free energy profile is similar to Q_10_ binding up to Q entrance in the BS (sdCOM<1.5 nm, Fig. 4A). The only difference is the relative smaller stability of the hydrated BS pose for Q_2_ (sdCOM=1.1 nm). Hydration of the Q_2_-tail and the chamber interior, and contacts performed by the Q_2_-head (Fig. 4B-D) are also equivalent to those performed by Q_10_ up to this point of the binding process.

**Figure 4:**
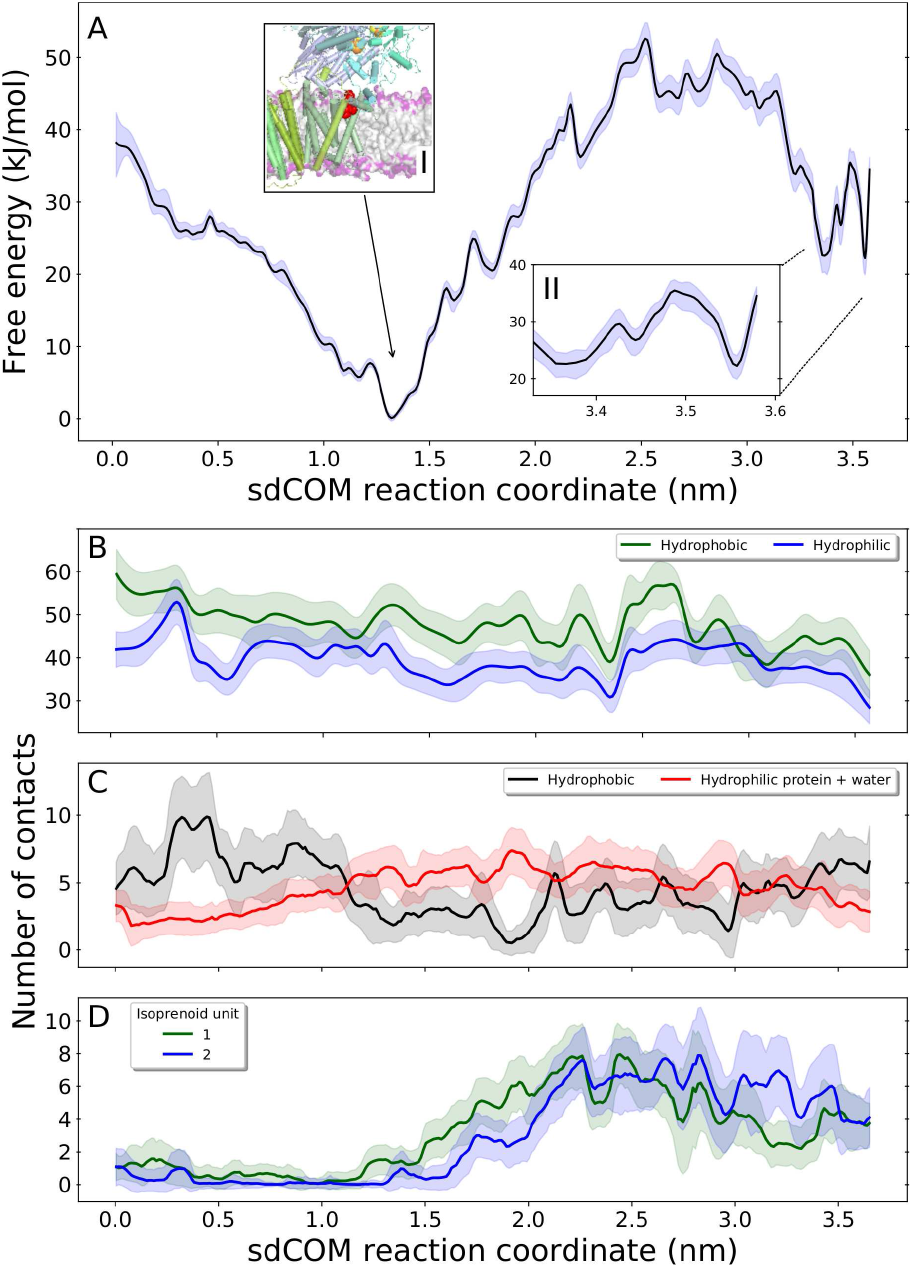
Binding of Q_2_ along the chamber in complex I. (A) Free energy profile (black line) with statistical uncertainty (blue shadow). Inset I shows the structure of complex I with Q_2_ at the binding site (BS) and inset II zooms in the RS region. (B) Water contacts to hydrophobic (green) and hydrophilic (blue) protein residues internal to the Q-chamber. (C) Q oxygen contacts with hydrophobic (black) and hydrophilic (red) groups; (D) Water contacts with Q_2_ isoprenoid units 1 (green) and 2 (blue).

As Q_2_ proceeds into the chamber, the free energy increases steeply and reaches a rough plateau of ~45 kJ/mol between 2.3<sdCOM<3.1 nm (Fig. S3C). Unfavorable hydration of the isoprenoid tail and of hydrophobic residues inside the Q-chamber also increase in this profile region (Figs. 4B and D).

At sdCOM~3.2 nm the free energy decreases simultaneously to dehydration of the Q_2_-tail and of chamber residues near the more hydrophobic RS (Figs. 4D and S6D). During the Q_2_ binding process, water contacts with hydrophobic residues inside the chamber decrease for sdCOM≤1.7 nm when the Q-tail is occupying the BS, then increase until sdCOM~2.7 nm due to re-hydration of residues near the BS and the chamber exit, and decrease again as the Q-head approaches the RS (Figs. 4B and S6E).

The barrier at sdCOM~3.4-3.5 nm separating the RS is independent of Q-tail length (13 kJ/mol for Q_2_ and 15 kJ/mol for Q_10_, Figs. 4A and 2A) and corresponds to dissociation of the hydrogen bond formed between the twisted His38 side chain and a Q carbonyl oxygen. As the Q-head enters the RS, His38 flips back and reforms contacts with the Q-head and Nqo4 D139 (Figs. 3B and S7E). At the RS, Q_2_ has only one clear local free energy minimum (Fig. S5E) which is 20 kJ/mol more stable than the Q_10_ minimum at the RS.

## Discussion

### Binding stability depends on Q-head interactions and on counterbalanced hydration of Q-tail and chamber residues

The free energy profiles estimated here for Q binding into complex I may be rationalized by the interplay of Q-head interactions and hydration of both Q-tail and apolar residues inside the amphipathic Q-chamber.

For initial binding of either Q_2_ or Q_10_, the Q-head exchanges unfavorable hydrophobic interactions with lipids and the apolar protein exterior for stabilizing hydrophilic interactions inside the BS. Hydration of the iso prenoid tail is minimal and entrance of Q expels to the aqueous phase part of the water molecules in contact with apolar residues near the chamber exit. These interactions and the considerable driving force (stabilization of 20-25 kJ/mol) for initial Q binding are independent of tail length.

However, internal hydration varies with the length of the Q-tail as binding proceeds towards the RS (Fig. S4). For Q_10_, hydrophobic residues in the chamber are continuously dehydrated due to internal water expulsion by the long Q-tail and the free energy increases mostly due to hydration of isoprenoid units. For Q_2_, both isoprenoid tail *and* apolar residues near the chamber exit are hydrated, leading to an even higher free energy. As Q_2_ approaches the more hydrophobic RS, the free energy decreases due to partial dehydration of the short isoprenoid tail. For Q_10_, the free energy remains relatively stable while approaching the RS because dehydration of primary isoprenoid units (1-2) is compensated by hydration of units (5-6) in the middle of the Q-tail. This balanced internal hydration is a particular case of hydrophobic effect that may be employed for selective recognition of amphiphilic ligands.^46,47^

The amphipathic nature of the Q-chamber (Fig. 1C) is conserved among species (Fig. S3D) suggesting the balance of interactions described here is essential for binding of the amphiphilic Q molecule. In fact, the role of long isoprenoid tails in dewetting the Q-chamber may have exerted part of the evolutionary pressure that led to complex I natural substrates with 6 or more isoprenoid units.

The extent of internal hydration in the Q-chamber may be questioned^48^ since the resolution of available complex I structures (3.3 Å at most^8,13^) is not enough to determine the position of water molecules. But, water access to binding sites and the interior of stable proteins is fast and frequently observed.^46,49,50^ The high water content found inside the Q-chamber here and in previous simulations of complex 1^24,51^ may also play a structural role for stabilization of the network of ionic residues found near the Q-chamber by shielding part of the unfavorable dielectric effect of charges buried in a protein.^52^ In mitochondrial complex I, external supernumerary units^9–11^ may decrease water penetration in the Q-chamber, and hence lead to a relatively less stable minimum for Q binding in the BS and lower free energies for Q reaching the RS.

### Mechanistic interpretations in comparison to previous experiments and future proposals

The high affinity BS found near the Q-chamber exit will be often loaded with a substrate molecule, even if Q is depleted from the membrane pool. Unwanted or reverse redox reactions involving Q bound in the BS will be minimized given the large distance to FeS centers.^53^

A sequential and separated BS in the Q-chamber^3,54^ is supported experimentally by the slow-relaxing EPR signal (SQ_Ns_) attributed to semiquinone radical interaction with cluster N2 separated by 30Å (Fig. 2B),^55–57^ and by auto-inhibition of complex I reductase activity observed in high concentrations of short-chain Q substrates.^42,43,58^

However, neither Q with a short or long tail have been observed bound to the BS in reported complex I structures. The intensive treatment with detergent used for protein purification will wash out endogenous Q. The BS has a broad free energy basin, where at least two Q-head poses were found (Fig. 3A). This will blur Q densities and difficult the resolution of a unique binding position. Dehydration of the water rich BS during crystallization may also destabilize Q binding at this site, in favor of the less hydrated RS. The relative free energy difference between both sites is quite low for short Q-tail and may depend on medium composition or even be reversed in crystals.^8,9^

Structural determination carried out using lipid phases with high content of Q_10_ or long-tail analogues may be able to observe Q bound to the BS. During enzyme turnover, Q binding to the BS is expected to involve a fast pre-equilibrium in relation to slower formation of the RS state. This could be probed by pre-steady state experiments of burst kinetics, analyzing the relaxation time in different substrate concentrations.^59^

The pre-RS pose (sdCOM=3.3 nm) found here with Q protonation only by His38 for both Q_10_ and Q_2_ suggests a secondary site for Q reduction and possible production of QH^(^·) semiquinone radicals.^56,57^ This could be an alternative reduction mechanism when re-protonation of Tyr87 during turnover has been impaired or when Tyr87 is mutated (Table S5).^60^ For two-electron reduction, the high basicity of Q^2-^ or even QH^−^ suggests that the second proton to form the quinol could be borrowed from a water molecule present in the chamber. The pre-RS site corresponds to the position occupied by the inhibitor 4-quinazolinylamine co-crystalized with the *Y. lipolytica* complex I structure.^9^ This site distance to the N2 cluster is 13-15 Å (Fig. 2B), also in line with the fast-relaxing EPR signal (SQ_Nf_) attributed to semiquinone formation.^55–57^

The experimental turnover rate of complex I catalysis is similar for reduction of Q with 10 to 4 isoprenoid units (average *k_cat_*=380±39 s^−1^), but decreases significantly for Q with shorter tail (*k_cat_*=138±7 s^−1^ for Q_2_ reduction).^48^ This rate drop-off may be interpreted by assuming the intrinsic barrier for redox conversion of oxidized Q to quinol (QH_2_) bound at the RS is independent of Q-tail length and by noting that free energies calculated here show similar barrier heights for Q_2_ and Q_10_ but higher stability for Q_2_ (favored by 20 kJ/mol when bound at or near the RS). Thus, the rate-limiting step in the slower Q_2_ turnover should change in relation to Q with long tails and take place after formation of bound quinol, probably product release. Due to high stability of Q bound to the BS and possible fast Q entrance inside complex I (high *k_on_*), a second short-tail substrate molecule may easily enter the chamber after the first one, and block dissociation once the first is reduced. This would not occur for Q with a long tail which occupies the BS and is also in line with the auto-inhibition of complex I activity observed in high concentrations of Q with short tails.^42,43,58^

Another molecular dynamics simulation study of Q binding into the chamber in complex I was recently published.^21^ A free energy profile obtained with umbrella sampling was shown for Q_1_ binding and a stable BS was also found. The profile is qualitatively similar to the Q_2_ profile presented here, but with lower free energies for Q moving towards the RS. Q_10_ binding was also studied, with an approximate diffusion model. The resulting free energy profile is almost flat and completely different from the Q_10_ profile presented here. These differences may be due to the unpublished force-field, truncated complex I structural model and significantly shorter (10 × to 20 × less than here) simulation times used and precluded analysis of chamber hydration along the binding process.^21^

## Conclusions

Free energy profiles obtained with large-scale molecular dynamics simulations for binding of amphiphilic molecules Q_2_ and Q_10_ into complex I were presented here. A secondary reactive site was found near cluster N2 with Q-head protonation by the rotated His38 side-chain. This site may lead to an alternative Q redox chemistry. A binding site near the chamber exit was also found with two binding poses stabilized by hydrophilic Q-head contacts. The profiles also help to interpret previous complex I observables, propose novel experiments to confirm the stability of Q at the secondary binding site, and suggest that mechanistic inferences of complex I binding and activity of natural Q substrates should be made with caution when based on experiments with short tail analogues.

Interestingly, the energetics of Q transit inside the chamber towards the reactive site is determined by the balanced hydration of Q-tail and of apolar residues within the Q-chamber. The Q-tail length modulates substrate selectivity by shifting this balance: a long tail will expel more water molecules from the Q-chamber but also bear more isoprenoid groups hydrated by the remaining internal water, whereas a short tail will leave more water inside the chamber and in contact with hydrophobic residues. This mechanism may be employed by amphiphilic molecules binding to internal hydrated cavities in other membrane proteins, such as lipid flippases and scramblases^61^ or fatty acid-binding proteins.^62^

## Acknowledgements

We thank Vanesa V. Galassi (USP) for assistance with the initial model construction, Sandro R. Marana (USP) for fruitful discussions, Leonid Sazanov (IST, Austria) for showing us the revised *T. Thermophilus* structure before publication and several comments from participants of the GRC Bioenergetics Meeting in 2017 when an initial version of this work was presented. Funding from FAPESP (projects 14/21900-2 and 16/24096-5) and computational resources from the SDumont cluster in the National Laboratory for Scientific Computing (LNCC/MCTI) are gratefully acknowledged.

## Electronic Supplementary Information

Additional description of the computational methods, results & discussion, including seven figures, five tables, one animated video and additional references (^63–68^) are available online.

## Electronic Supplementary Information

### Supplementary Methods

The structure of complex I from *T. thermophilus*^8^ did not contained a bound NADH, so this substrate was not included in the computational model. The NADH binding site is located more than 50Å from the Q-chamber. Given the total simulation time per sampling window, it is unlike that any structural perturbation near the Q-chamber could have been caused by lack of NADH in the model. A disulfide bond was included between residues Nqo2 C144 and C172. All missing atoms in chain terminal regions were not built in the model nor considered for analysis. The only missing terminus that might be located close to the Q-chamber is Nqo6 residues 1-15 (sequence MALKDLFERDVQELE). This N-terminus has amphipathic character with many charged residues and the sequence is not conserved among species. Thus, it should not penetrate in the membrane nor perturb Q attachment to the nearby chamber exit formed by Nqo8 helices TM1, TM6 and AH1. It should be mentioned that other Q-chamber exits exposed to the lipid membrane have been proposed, including TM1 and TM7, or TM5 and TM6, both together with AH1, all from subunit Nqo8 (or the equivalent ND1 in bovine).^8,10,13^ These other proposals were not considered here as they show narrower exits and should require significant protein movement for passage of the Q-head.

Although Nqo4 loop β 1-β2 and Nqo8 loop TM5–TM6 have been observed in different conformations,^9–11^ the position of these loops in the *T. thermophilus* structure should be appropriate for describing Q binding as it was co-crystalized with Q analogues.^8^

All protein hydrogens atoms were built with the GROMACS utility^32^ during model construction. The NPT ensemble was used in all simulations. Temperature was maintained with the Bussi thermostat^63^ and a coupling constant of 0.1 ps with two separate coupling groups (water plus ions and everything else). Pressure was kept with the Berendsen barostat^64^ during the initial equilibration and with the Parrinello-Rahman barostat for the umbrella sampling simulations with a coupling constant of 1 ps and a compressibility of 1.10^−5^ bar^−1^. Semi-isotropic coupling in the direction normal to the bilayer was applied. A real space cutoff of 1.2 nm was used for PME electrostatics and van der Waals interactions. No dispersion corrections were applied. All covalent bonds were constrained using LINCS.^65^

For both Q_10_ and Q_2_ simulations, US windows were separated by 0.1 nm in the dCOM range 0.43-3.43 nm and 3.60-4.10 nm that covers binding through the entire Q-chamber and the membrane exit. Residues used for the definition of both reaction coordinates are shown in Fig. S1. Additional windows were placed with reference dCOM and Path CV as shown in Table S1. Coordinate distributions were checked for sufficient overlap between adjacent 2D-windows, as required by the WHAM procedure. Two Supplementary Files are included: coordinates of the 30 milestone configurations used for calculation of Path CV; and the initial Q_10_-bound structure used for all simulations. Results can be fully reproduced using these files and the information described here. In all analysis shown, a contact was considered when the distance between any pair of atoms in each group was smaller than 3.5Å. Table S2 shows the atoms included on each group. Isoprenoid units were composed of 5 carbons and the 8 bound hydrogens, except for the terminal isoprenoid-10 with an extra hydrogen.

The choice of reaction coordinate is crucial for reliable calculation of free energy profiles. A simple distance metric between a reference point in the protein and another in the ligand would suffice to describe transit along the Q-chamber if it was homogeneously thin and tight, like a perfect tunnel. However, the Q-chamber BS and RS are relative bulky cavities where different poses and interactions may correspond to the same reference distance. In these sites, a simple distance metric becomes ill-defined and sampling with only one control coordinate such as dCOM would be flawed. In order to help resolving Q transit inside the full chamber, the Path CV coordinate based on milestone configurations along the binding pathway was used here.^17^ Two-dimensional free energy surfaces are shown in Fig. S5A and B. They are rather diagonal, indicating that expression of results using projections on either dCOM or Path CV coordinates would lead to equivalent conclusions. For instance, the projected profiles in Fig. S5C and D are qualitatively similar displaying the same number of local minima, maxima and relative energies, except for the end of the Path CV profile (Path CV>26) which is more compressed due to the smaller number of milestone configurations used in this region. Two-dimensional profiles are used here to resolve the profile minima in the RS region, as shown in Figs. 3C and Fig. S5E.

Given the size and complexity of the simulated system, it is unlike that all degrees of freedom were fully sampled in our simulations. However, the most important ones – hydration of the chamber interior and Q-tail – appear to have been well sampled. Water exchange from the protein interior was fast (~2 ns)^50^ and hydration of different groups in the Q-chamber and the Q-tail equilibrated in less 50 ns in all US windows. Figs. S2D/E show that stabilization within statistical uncertainty of the free energy calculated for Q bound in the RS state *(ΔG*_RS_) is observed with an equilibration time of 40-50 ns and acquisition time of 80-90 ns per US window. This point of the profile has the slowest convergence over all US simulations presented here. Error bars (blue shadows) in free energy profiles indicate the sampling statistical uncertainty only. Imperfections in model composition and structure or in the interaction force-field may also introduce systematic errors, which unfortunately are difficult to estimate. Nevertheless, the favorable comparison between three different profiles obtained from different starting structures for Q_10_ (Figs. 2A and S2C) and Q_2_ (Fig. 4A) suggest that our results have at least semi-quantitative precision.

### Supplementary Results & Discussion

#### Structure of Nqo6 loop *α*_2_–*β*_2_

As described in the Methods section of the main text, the protein model used here was based on the structure of the respiratory complex I from *T. thermophilus* (PDB ID: 4HEA)^8^, with Nqo6 loop *α2–β2* rebuilt using the loop in the ovine enzyme as a template (Fig. S2A). The resulting loop structure after relaxation is very close to other mammalian enzymes, including the recent mouse structure (PDB ID: 6G2J)^13^ determined with a 3.3Å resolution (RMSD of 0.14 nm, Table S3). The original (4HEA) Nqo6 loop α2–β2 structure is quite different (RMSD of 0.56 nm). Hence the RMSD of 0.18 nm found for Q-chamber in comparison to the 4HEA structure in Table S3 is caused completely by deviations on this Nqo6 loop.

In preliminary simulations shown in Fig. S2, the missing Nqo6 residues 65-69 were built with M0DELLER *de novo, i.e.* without a structural template. A rather different loop structure was obtained as shown in Fig. S2B, with divergent positions of highly conserved Arg69 and Arg62. This difference had a profound impact on the calculated free energy (Fig. S2C). The profile in sdCOM<1.5 nm is similar to Fig. 2A in the main text, as Q is still distant from Nqo6 loop α2–β2. However, the bumps found in between 1.5<sdCOM<2.5 nm (Fig. S2C) are due to artificial interactions of Arg62 with the Q-head. These are not observed in the main text simulations as Arg62 is distant from the Q-chamber when the Nqo6 loop structure is based on the ovine enzyme (Fig. S2B). Differences in the profiles are even more pronounced when Q is bound at the RS. In fact, no (meta)stable free energy minimum is found in the RS, as Arg69 artificially binds the Q-head instead of the protonated Nqo4 His38 (Fig. S2B). Thus, caution should be used when building loop segments *de novo*, particularly if the built residues are in contact or near the protein region to be studied in detail.

#### Modelling limitations

Based on the complex I structure co-crystalized with a substrate analogue,^8^ our initial model should be appropriate to study Q binding in the internal chamber formed by subunits Nqo4, Nqo6 and Nqo8. This Q-chamber is rather stable and shows similar conformations among different species (Table S3). But the flexible Nqo6 loop α2–β2, originally with a missing segment,^8^ had its structure rebuilt here using the ovine loop^11^ as a template. Such structural refining turned out to be important to describe Q stability in the RS (Fig. S2).

Given an appropriate structural model and a reasonable description of Q interactions,^22^ accurate simulation of Q binding also requires sampling the configurational space. The narrow and conformationally restrict Q-chamber suggests that configurational freedom of groups inside will be small and that description of ligand binding with a simple distance coordinate may be accurate. However, the BS and the RS are bulky cavities and a more elaborate control coordinate such as the Path CV^17^ had to be employed (see SI Methods). Formation of the initial ligand-protein encounter complex with Q still in the membrane or crawling over the protein surface in order to find the exposed chamber exit were not considered here. Although these may contribute to the binding free energy, simulating processes with Q outside the chamber requires sampling a much larger configurational space or using elaborate spacial restrictions.^66^

#### Protein contacts and flexibility along Q-binding

Q transit may be described starting from the chamber exit when the Q-head is in contact with apolar residues in helices Nqo8 TM1, TM6 and AH1 (Fig. S3B). 0nce inside the chamber and towards the BS, the Q-head is close to the Nqo8 loop TM5-TM6 (in the membrane arm side) and TM1 (in the opposite side), and with Nqo6 β 1-strand and loop α1-β1. After the BS, when the chamber has a turn upwards, the Q-head interacts mostly with Nqo6 helix α2, loop α2–β1, β1-strand and turn α4-α5. 0nce in the RS, the Q-head forms contacts with Nqo6 *α2* (and the nearby loop α2–β1 that binds the N2 cluster) and Nqo4 loop β 1-β2, end of α4 and C-terminus helices and loop α2-α3 (Fig. 1B).

The Q-chamber backbone and all described secondary structures remain stable during the simulations. For Q_10_, root mean-squared deviations (RMSD) of *C_α_* position for chamber residues vary between 0.10-0.20 nm along the binding coordinate. Most of this can be traced to thermal fluctuations of Nqo4 loop β1-β2, Nqo6 loop α2–β2 and Nqo8 loop TM5-TM6 (Table S3).

Besides the BS, pre-RS and RS positions already described in the main text, the Q-head may directly contact charged residues at sdCOM=1.8 nm and hydrogen bond with Nqo8 Arg45 and Nqo6 Arg69 (Fig. S3A). The Q-tail forms several hydrophobic contacts with apolar residues particularly when isoprenoid units approach the chamber exit, the Nqo6 helix α2 and the Nqo4 loop β 1-β2 (Fig. 1).

#### Point mutations and Q-chamber stability

Point mutations on residues in the Q-chamber perturb assembly of the complex and its enzymatic activity (Table S5).^8^ Several of these mutations are known to be pathological in humans.^6,7^ Interestingly, mutations on the less flexible residues Nqo8 E35, R36, R45, D62, performing stable ionic contacts (Fig. S7) in the opposite side of the membrane arm, near Nqo8 helices TM1 and AH1, are more deleterious and reduce the content of assembled complex I, unless a conservative mutation is observed. These four residues also show the highest conservations among the Q-chamber (Table S4). There is more plasticity on the network of ionic contacts on the side of the E-channel which fluctuate often during our simulations (Fig. S7). Mutations (Nqo8 E163, E213, R216, E223, E225, E248 and R294) are less deleterious and activity is maintained even when a charge is reversed (such as E163K, Table S5). Thus, ionic contacts on the side of Nqo8 helices TM1 and AH1 have an essential structural role and keep the Q-chamber architecture. Contacts on the E-channel side near Nqo8 helices TM5 and TM6, and Nqo6 β1 strand may fluctuate without disrupting the enzyme structure. These “controlled” fluctuations may be explored by evolution in order to modulate enzyme activity. Indeed, it has been shown recently that localized unfolding of Nqo8 TM5 and TM6 and Nqo6 β 1 is involved in the active-to-deactive transition observed in mammalian complex I.^67^

#### Possible role of Q binding in activation of the E-channel

The network of ionic contacts near the chamber is perturbed by passage of Q, specially contacts located on the membrane arm side (Fig. S7). The Q-head forms hydrogen bonds with Nqo8 Arg36 in the BS (sdCOM=1.4 nm, Fig. 2F), leading to rupture of the Arg36-Glu66(Nqo6) salt bridge, which otherwise remains stable along Q binding. Contact Arg36-Asp62, although formed transiently before the BS, is only stable in sdCOM>1.4 nm, after Q passage towards the RS (Fig. S7C). Nqo6 Arg69 forms transient salt bridges with both Nqo8 Glu223 and Glu225. After passage of Q and hydrogen bonding with the Q-head (sdCOM=2.3 nm, Fig. 2F), only Arg69 bridge with Glu225 remains stable and the contact with Glu223 may no longer be formed (sdCOM>2.3 nm, Fig. S7B). When not coordinated with Arg69, the flexible Glu223 approaches Nqo8 Arg216 (in sdCOM=0.3, 1.2, 2.2 and 3.1 nm, Fig. S7D) disturbing its contact with Glu213, which in turn may approach Glu163 (Fig. S7F), the following residue in the E-channel (Fig. 1B). Note that Glu223 never approaches Glu213 itself (Fig. S7E). Additionally, when Q is in the RS (sdCOM> 3.5 nm), the otherwise stable bridge Arg216-Glu248 in Nqo8 is slightly perturbed (Fig. S7D), bridge Arg216-Glu213 has the closest contact over the binding process and Glu213 is the farthest from Glu163 (Fig. S7F).

This highly hydrated network of ionic contacts has been proposed to be involved in a putative Nqo8 proton pumping channel and in the coupling between the Q redox process and proton pumping in the membrane arm, via the E-channel.^8,21,24,51^ The above observations allow us to speculate a possible role for Q transit in the coupling mechanism. When Q is unbound from complex I, the proton pumps should be deactivated as no electron transport will take part. This is in line with the large Glu213-Glu163 distances seen for sdCOM<2.0 nm. As Q loads the chamber, the ionic network is perturbed as described above, Glu213 approaches Glu163, and the E-channel may propagate a “ready-to-go” signal to the proton channels. As soon as Q is reduced and binds the H+ from Nqo4 His38, Asp139 (Asp160 in bovine numbering) may twist and contact Arg294 which in turn complete activation of the E-channel to trigger proton transfer.

In summary, the network of charged residues close to the BS may function as an ionic interface separating the hydrophobic exterior (apolar residues and lipid) from the more hydrated Q-chamber interior, much like the zwitterionic head in a phospholipid is at the membrane interface (Fig. 2B). Accurate molecular simulations of the correlated motions found on this network of residues in complex I may require more realistic model compositions such as representing the electrochemical gradient across the membrane.

**Figure S1:**
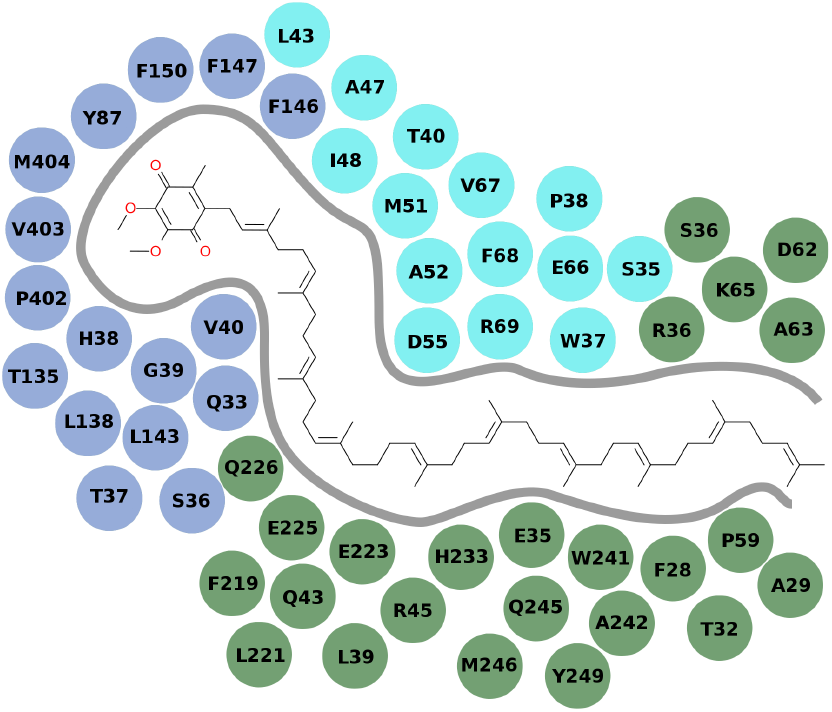
Residues along the Q-chamber which *C_α_* was used for definition of dCOM and Path CV reaction coordinates. Bound Q_10_ is also shown. For dCOM, Nqo4 (light blue) residues Q33, H38, G39, V40, T135, L143, T144, P402, V403 and M404 were used. For Path CV, Nqo4 residues Q33, S36, T37, H38, G39, V40, Y87, T135, L138, L143, F146, F147, F150, P402, V403, M404, Nqo6 (cyan) residues S35, W37, P38, T40, L43, A47, I48, M51, A52, D55, E66, V67, F68, R69, A70, and Nqo8 (pale green) residues F28, A29, T32, E35, R36, L39, Q43, R45, P59, D62, A63, K65, S66, F219, L221, E223, E225, Q226, H233, W241, A242, Q245, M246, Y249 were used.

**Figure S2:**
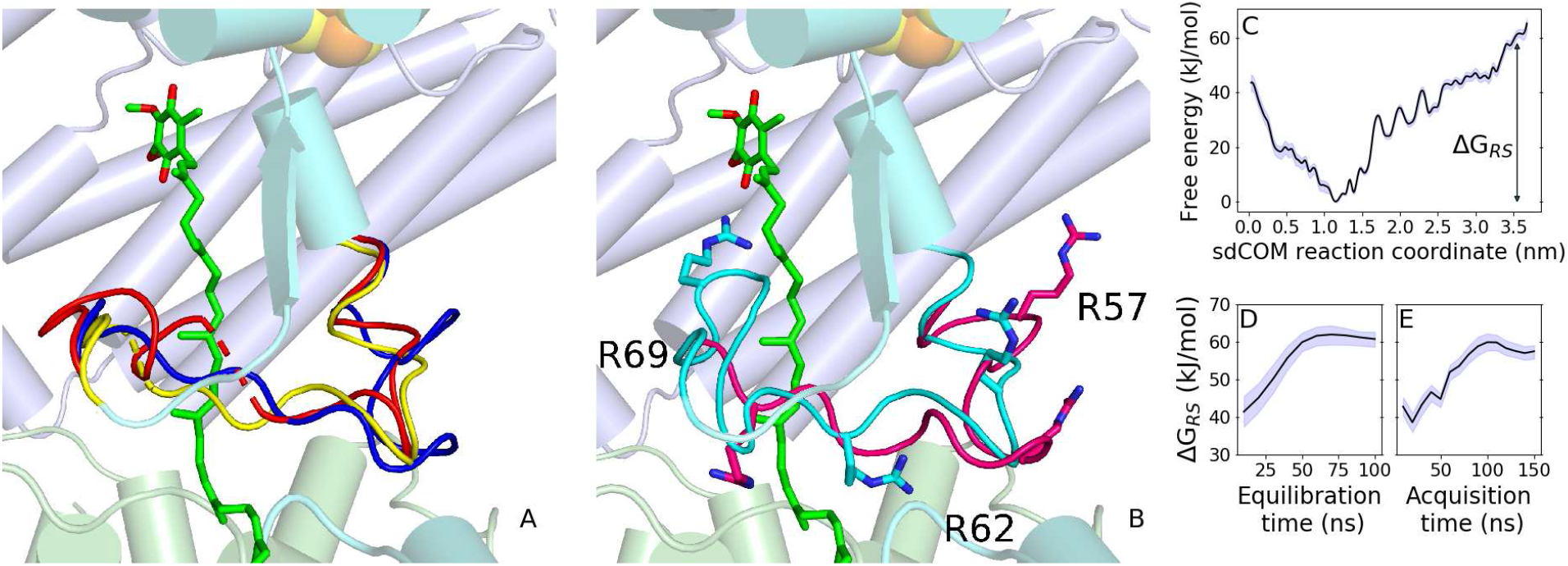
Structure of the Nqo6 loop α2–β2 and its impact on the free energy profile for Q binding. (A) Close view in cartoon of subunits Nqo4 (light blue), Nqo6 (cyan) and Nqo8 (pale green) with Q_10_ modeled in green sticks and loop structures from ovine in blue (PDB ID 5LC5), *T. thermophilus* in yellow (PDB ID 4HEA) and *Y.lipolytica* in red (PDB ID 4WZ7). Note the missing segment 65-69 from the *T. thermophilus* in dashed yellow backbone. (B) Same view as (A), but with the loop of the relaxed initial protein model used in the main text simulations (Fig. 2) in magenta (compare with the blue loop in panel A), and the loop from *T. thermophilus* with the missing residues 65-69 built with MODELLER *de novo* in cyan. Arg69, Arg62 and Arg57 are also shown. (C) Free energy profile in black with statistical uncertainty in blue shadow obtained with the Nqo6 segment 65-69 built *de novo* as shown in panel B. Free energy convergence of Q bound in the RS (*ΔG_RS_*) with increasing (D) equilibration and (E) acquisition simulation time per US window.

**Figure S3:**
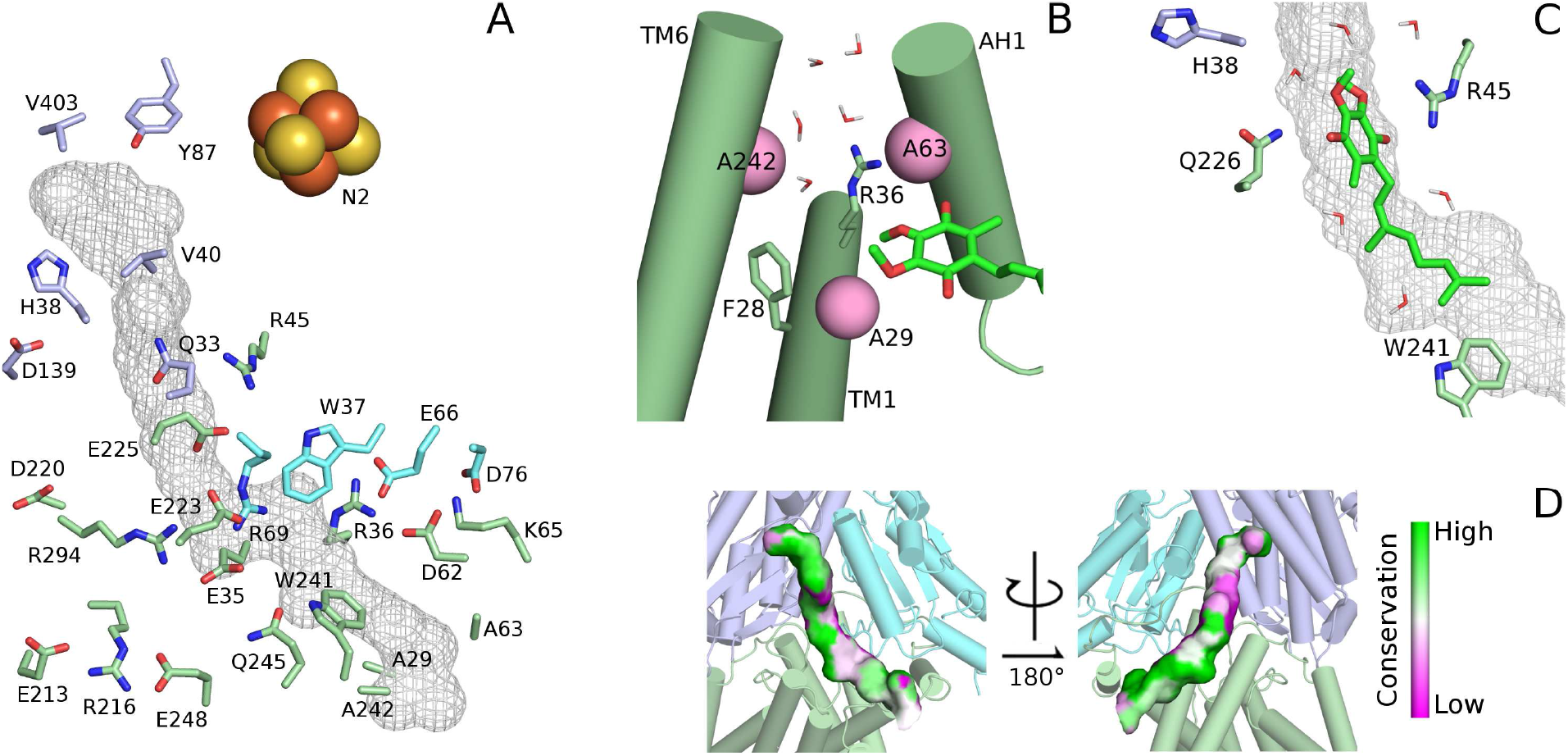
Protein environment of Q in the binding chamber. Key residues are shown for subunits Nqo4 (in light blue), Nqo6 (cyan) and Nqo8 (pale green). (A) Chamber overview with residue position and molecular surface mesh of Q bound in the RS. (B) CE position, with Q outside the chamber exit. Helices TM1, TM6 and AH1 are shown in cartoon with Ala residues em pink spheres. Water molecules are shown to indicate chamber hydration. This is the only panel displayed in a different viewing perspective. (C) Q_2_ in green on the high free energy region for its binding (sdCOM=2.5 nm). Surface mesh as in panel A. (D) Molecular surface of Q bound in the RS colored by conservation of the nearest protein residue in *T. thermophilus* structure. The conservation score was calculated as described in Table S4.

**Figure S4:**
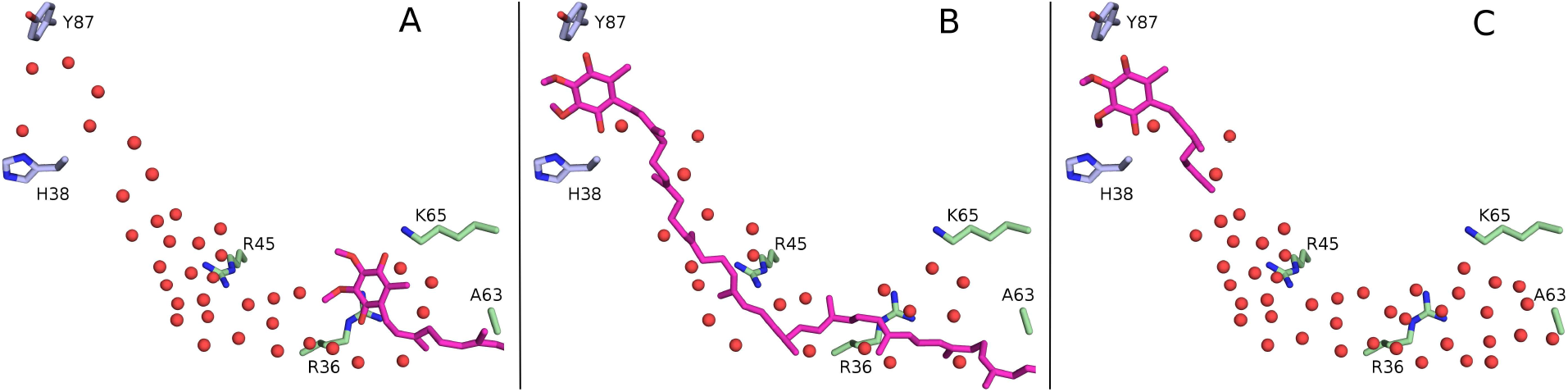
Chamber internal hydration with water oxygens shown in red spheres. Q in magenta occupies the (A) BS, (B) Q_10_ in RS, and (C) Q_2_ in RS positions.

**Figure S5:**
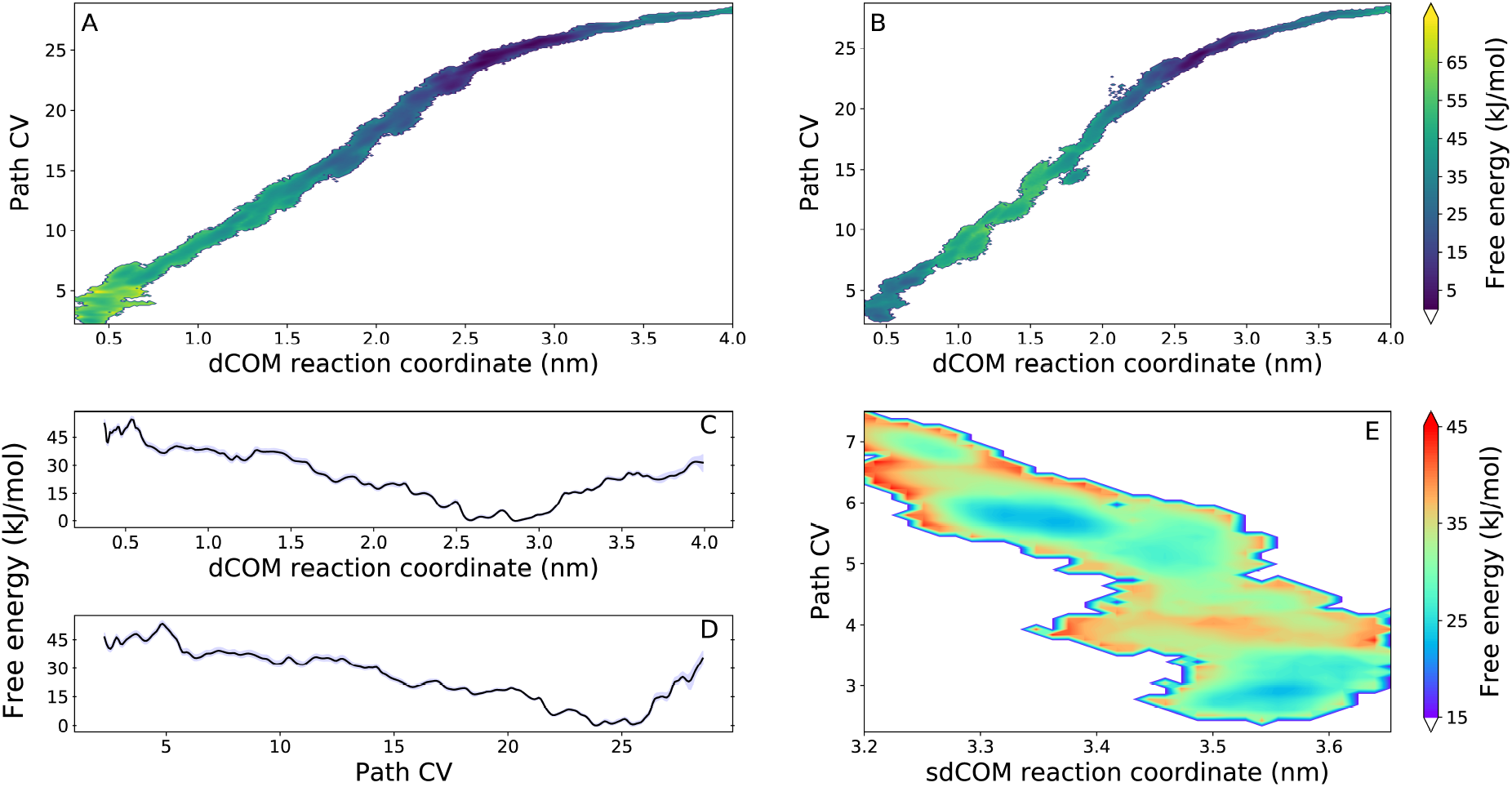
Free energy profiles for Q binding along the chamber in complex I. Complete twodimensional free energy profiles calculated for (A) Q_10_ and (B) Q_2_ binding. Projection of the Qio 2D minimum free energy on (C) dCOM and (D) Path CV coordinates. Note the dCOM coordinate (instead of the shifted sdCOM) is used in the x-axis of panels A-D. Here, binding from the membrane into the chamber proceeds from high to low values for dCOM and Path CV reaction coordinates. (E) Two-dimensional free energy profile for Q_2_ binding in the RS region.

**Figure S6:**
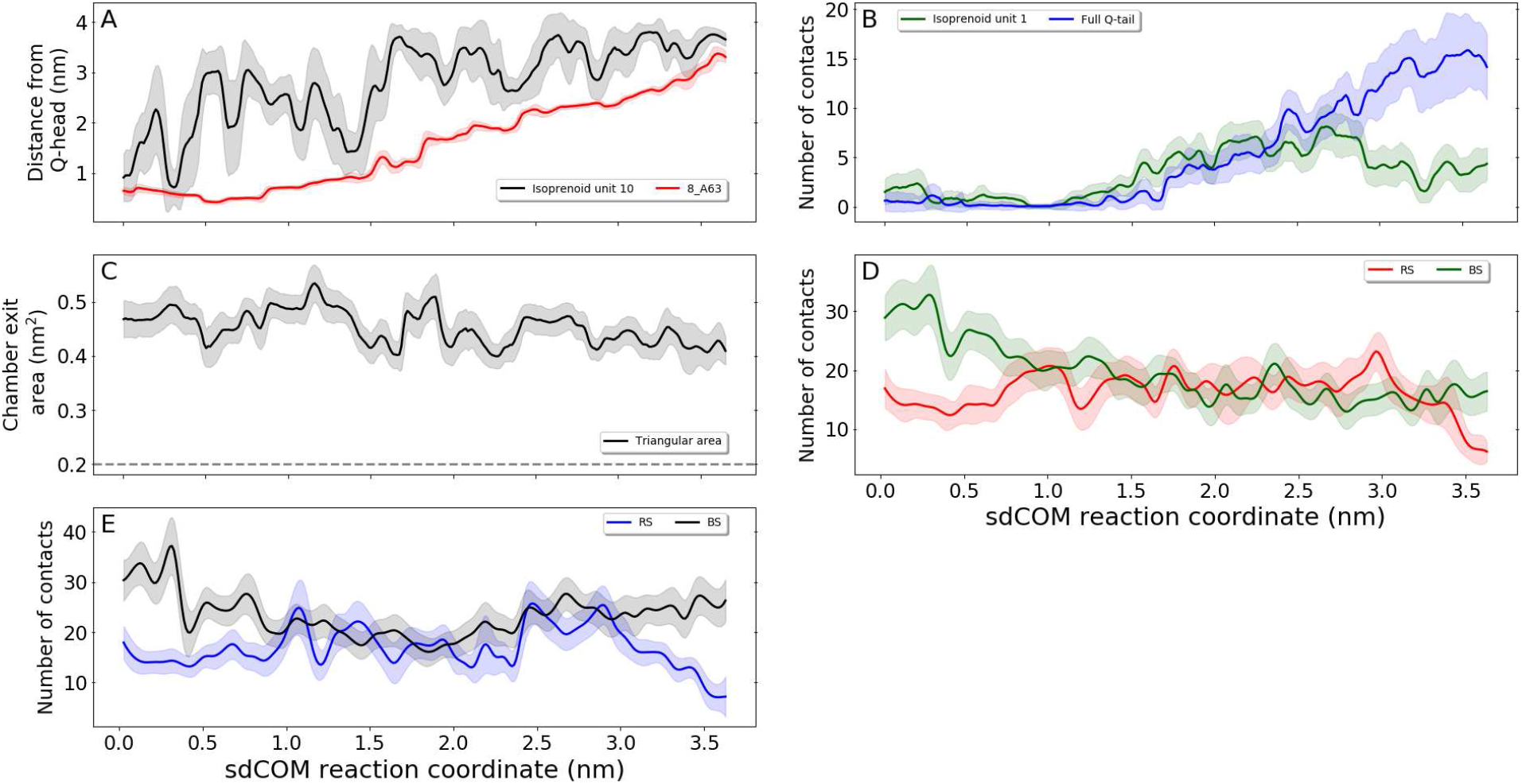
Additional simulation analysis. (A) Distance of the Q-head COM to the C_α_ from Nqo8 Ala63 in the chamber exit in red and to the COM of isoprenoid unit 10 in black. (B) Hydration of the full Q_10_-tail in blue and of isoprenoid unit 1 in green. (C) Area of the triangle determined by the C_α_ position of Nqo8 Ala29 (helix TM1), Ala63 (helix AH1) and Ala242 (helix TM6) in the Q-chamber exit to the lipid membrane (Fig. S3B). For the original crystal structure (PDB 4HEA),^8^ this area is 0.37 nm^2^. The dashed gray line shows the estimated transversal area of the Q-head (0.2 nm^2^=0.7 nm × 0.3 nm, respectively the distance between carbonyl oxygens and the van der Walls thickness of the Q ring). Total hydration of BS and RS residues in the Q-chamber (D) for Q_10_ and (E) for Q_2_ binding.

**Figure S7:**
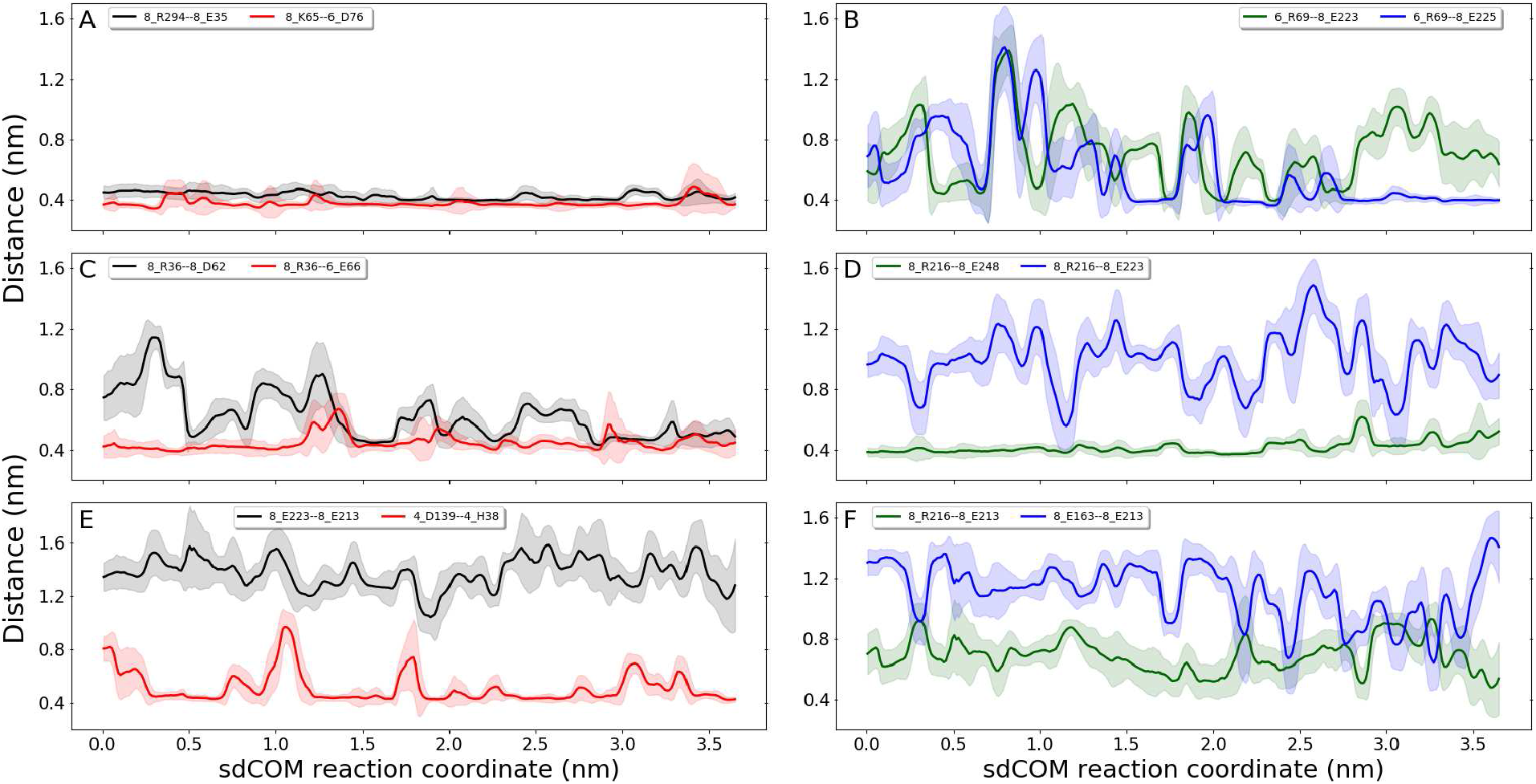
Network of ionic contacts near the Q-chamber. Pair distances along Q binding between side chains of (A) Nqo8_R294-8_E35 (black) and 8_K65-6_D76 (red), (B) 6_R69-8_E223 (green) and 6_R69-8_E225 (blue), (C) 8_R36-8_D62 (black) and 8_R36-6_E66 (red), (D) 8_R216-8_E248 (green) and 8_R216-8_E223 (blue), (E) 8_E223-8_E213 (black) and 4.D139–4_H38 (red), and (F) 8_R216-8_E213 (green) and 8_E163-8_E213 (blue). Distances were computed between C_ς_ for Arg, N_ς_ for Lys, C_δ_ for Glu, C_γ_ for Asp and COM of imidazole ring for His.

**Table S1:** Reference dCOM (in nm) and Path CV of additional windows used in US simulations^a^.

| Q<sub>10</sub> |  | Q<sub>2</sub> |  |
| --- | --- | --- | --- |
| dCOM | Path CV | dCOM | Path CV |
| 0.42 | 3.31 | 0.48 |  |
| 0.43 | 2.80 | 0.48 | 3.90 |
| 0.43 | 3.10 | 0.58 | 5.60 |
| 0.43 | 3.20 | 0.73 | 6.40 |
| 0.43 | 3.50 | 1.11 | 10.0 |
| 0.46 | 3.72 | 1.85 | 16.7 |
| 0.48 |  | 2.33 | 21.8 |
| 0.48 | 3.90 | 2.77 |  |
| 0.48 | 4.30 | 3.52 | 27.6 |
| 0.48 | 4.60 |  |  |
| 0.48 | 4.90 |  |  |
| 0.54 | 5.10 |  |  |
| 0.54 | 5.20 |  |  |
| 1.43 | 12.0 |  |  |
| 1.93 | 16.6 |  |  |
| 2.48 |  |  |  |
| 2.77 |  |  |  |
| 3.20 | 26.5 |  |  |

<sup>a</sup> In addition to windows in dCOM range 0.43-3.43 nm and 3.60-4.10 nm separated by 0.1nm. No restrain was applied to Path CV when a reference value for this coordinate is not listed.

**Table S2:** List of atoms included on each group used for simulation and analysis.

| Group | Atoms |
| --- | --- |
| Water | Oxygen (OW) |
| Q-head | Heavy atoms in Q except for the isoprenoid tail |
| Q-head oxygens | Four oxygen atoms in Q |
| Lipid tail | All atoms in both palmitoyl and oleoyl POPC hydrocarbon chains (except for carbonyl group) |
| Q-chamber C<sub>α</sub> | C<sub>α</sub> of residues Nqo4 33, 37, 38, 39, 40, 62, 87, 88, 131, 132, 135, 138, 143, 146, 147, 150, 398, 401, 402, 403, 404, residues Nqo6 37, 40, 41, 42, 43, 44, 45, 47, 48, 51, 52, 67, 68, 69, 70, 81 and residues Nqo8 28,29, 32, 35, 36, 39, 45, 59, 60, 62, 63, 65, 216, 219, 225, 226, 241, 242, 245, 246, 249, 294. |
| Nqo4 loop $\beta 1-\beta 2$ | Nqo4 residues 32-41 |
| Nqo6 loop $\alpha 2-\beta 2$ | Nqo6 residues 55-70 |
| Nqo8 loop TM5-TM6 | Nqo8 residues 220-238 |
| BS residues | Side-chains of residues Nqo8 28, 29, 32, 35, 36, 59, 60, 62, 63, 65, 226, 241, 242, 245, 246, 249, 294 |
| RS residues | Side-chains of residues Nqo4 38, 39, 87, 88, 131, 132, 135, 150, 398, 401-404 and Nqo6 44, 45, 48 |
| Apolar (hydrophobic) protein | Side chains of G, P, A, V, I, L, M, F and W (except for N<sub>ε1</sub> and H<sub>ε1</sub>) |
| Charged and polar (hydrophilic) protein | Backbone N and hetero atoms in side chains of R, K, D, E, H, S, T, N, Q, C, Y |

**Table S3:** Root mean-squared deviation (in nm) of the C_α_ position for different structural models and groups of residues in the Q-chamber. The initial protein model (see Methods) was used as a reference structure.

| Group | Structural model<sup>a</sup> |  |  |  |  |  |
| --- | --- | --- | --- | --- | --- | --- |
|  | Q<sub>10</sub> simulation | 4HEA | 4WZ7 | 5LC5 | 5LNK | 6G2J |
| Q-chamber | 0.10-0.20 | 0.18 | 0.24 | 0.17 | 0.21 | 0.16 |
| Nqo4 loop $\beta 1-\beta 2$ | 0.12-0.26 | 0.02 | 0.25 | 0.21 | 0.38 | 0.20 |
| Nqo6 loop $\alpha 2-\beta 2$ | 0.13-0.21 | 0.56 | 0.47 | 0.18 | 0.15 | 0.14 |
| Nqo8 loop TM5-TM6 | 0.14-0.29 | 0.03 | 0.55 | 0.24 | 0.47 | 0.24 |

<sup>a</sup> The average plus one-standard deviation RMSD range observed over the full reaction coordinate of the Q<sub>10</sub> US simulation is shown. The given structural codes correspond to PDB IDs 4HEA for *T. thermophilus*,<sup>8</sup> 4WZ7 for *Y. lipolytica*,<sup>9</sup> 5LC5 for bovine,<sup>10</sup> 5LNK for ovine<sup>10</sup> and 6G2J for mouse.<sup>13</sup>

**Table S4:**
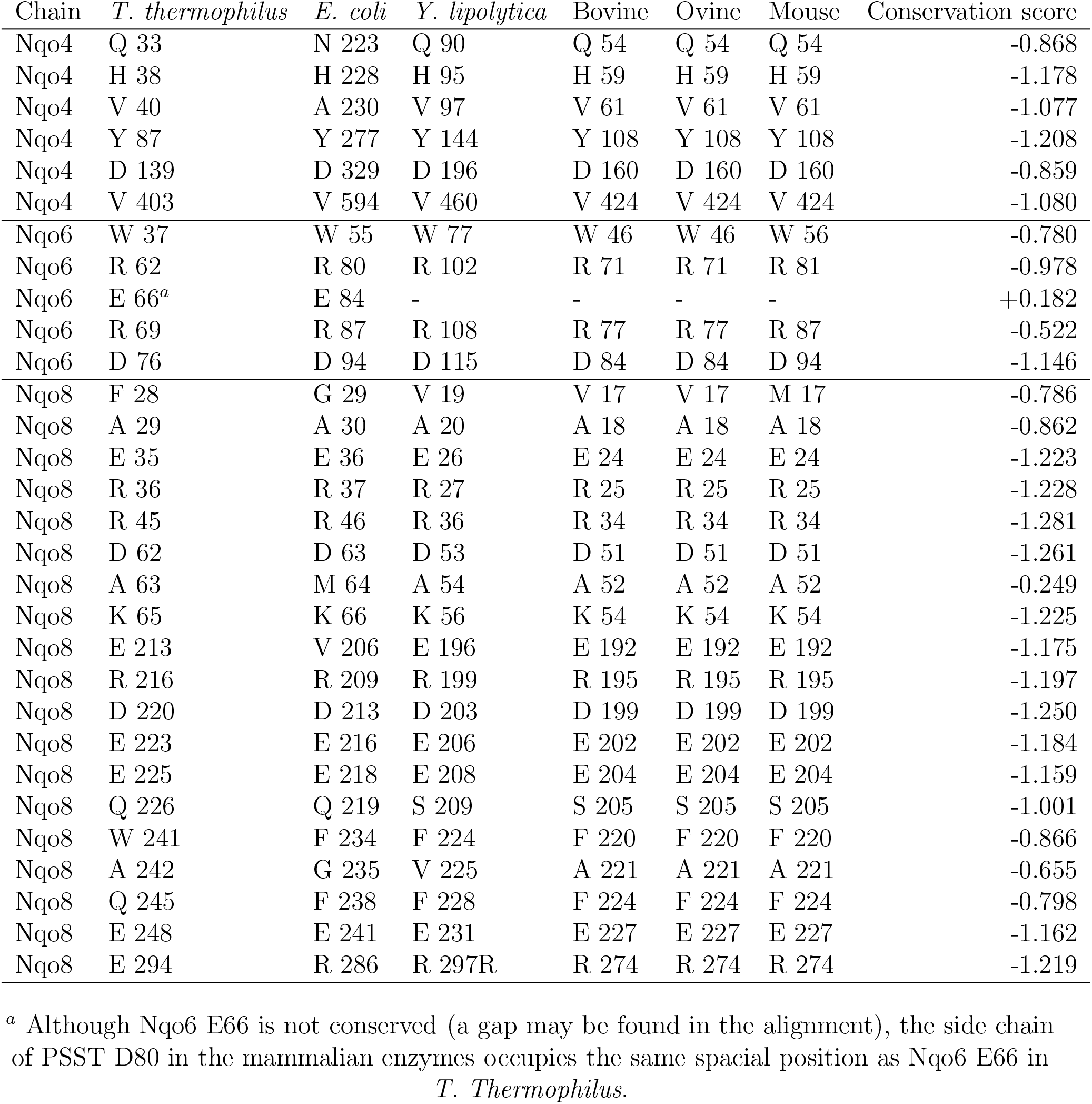
Conservation among species of Q-chamber residues in subunits Nqo4, Nqo6, Nqo8. Score was calculated after a multiple sequence alignment of 300 species with ConSurf 2016^68^ (using a maximal identity between sequences of 90% and a minimal identity for homologs of 30% as elsewhere,^8^ with otherwise default settings). More negative scores indicate higher conservation.

| Chain | *T. thermophilus* | *E. coli* | *Y. lipolytica* | Bovine | Ovine | Mouse | Conservation score |
| --- | --- | --- | --- | --- | --- | --- | --- |
| Nqo4 | Q 33 | N 223 | Q 90 | Q 54 | Q 54 | Q 54 | -0.868 |
| Nqo4 | H 38 | H 228 | H 95 | H 59 | H 59 | H 59 | -1.178 |
| Nqo4 | V 40 | A 230 | V 97 | V 61 | V 61 | V 61 | -1.077 |
| Nqo4 | Y 87 | Y 277 | Y 144 | Y 108 | Y 108 | Y 108 | -1.208 |
| Nqo4 | D 139 | D 329 | D 196 | D 160 | D 160 | D 160 | -0.859 |
| Nqo4 | V 403 | V 594 | V 460 | V 424 | V 424 | V 424 | -1.080 |
| Nqo6 | W 37 | W 55 | W 77 | W 46 | W 46 | W 56 | -0.780 |
| Nqo6 | R 62 | R 80 | R 102 | R 71 | R 71 | R 81 | -0.978 |
| Nqo6 | E 66<sup>a</sup> | E 84 | - | - | - | - | +0.182 |
| Nqo6 | R 69 | R 87 | R 108 | R 77 | R 77 | R 87 | -0.522 |
| Nqo6 | D 76 | D 94 | D 115 | D 84 | D 84 | D 94 | -1.146 |
| Nqo8 | F 28 | G 29 | V 19 | V 17 | V 17 | M 17 | -0.786 |
| Nqo8 | A 29 | A 30 | A 20 | A 18 | A 18 | A 18 | -0.862 |
| Nqo8 | E 35 | E 36 | E 26 | E 24 | E 24 | E 24 | -1.223 |
| Nqo8 | R 36 | R 37 | R 27 | R 25 | R 25 | R 25 | -1.228 |
| Nqo8 | R 45 | R 46 | R 36 | R 34 | R 34 | R 34 | -1.281 |
| Nqo8 | D 62 | D 63 | D 53 | D 51 | D 51 | D 51 | -1.261 |
| Nqo8 | A 63 | M 64 | A 54 | A 52 | A 52 | A 52 | -0.249 |
| Nqo8 | K 65 | K 66 | K 56 | K 54 | K 54 | K 54 | -1.225 |
| Nqo8 | E 213 | V 206 | E 196 | E 192 | E 192 | E 192 | -1.175 |
| Nqo8 | R 216 | R 209 | R 199 | R 195 | R 195 | R 195 | -1.197 |
| Nqo8 | D 220 | D 213 | D 203 | D 199 | D 199 | D 199 | -1.250 |
| Nqo8 | E 223 | E 216 | E 206 | E 202 | E 202 | E 202 | -1.184 |
| Nqo8 | E 225 | E 218 | E 208 | E 204 | E 204 | E 204 | -1.159 |
| Nqo8 | Q 226 | Q 219 | S 209 | S 205 | S 205 | S 205 | -1.001 |
| Nqo8 | W 241 | F 234 | F 224 | F 220 | F 220 | F 220 | -0.866 |
| Nqo8 | A 242 | G 235 | V 225 | A 221 | A 221 | A 221 | -0.655 |
| Nqo8 | Q 245 | F 238 | F 228 | F 224 | F 224 | F 224 | -0.798 |
| Nqo8 | E 248 | E 241 | E 231 | E 227 | E 227 | E 227 | -1.162 |
| Nqo8 | E 294 | R 286 | R 297R | R 274 | R 274 | R 274 | -1.219 |

<sup>a</sup> Although Nqo6 E66 is not conserved (a gap may be found in the alignment), the side chain of PSST D80 in the mammalian enzymes occupies the same spacial position as Nqo6 E66 in *T. Thermophilus*.

**Table S5:**
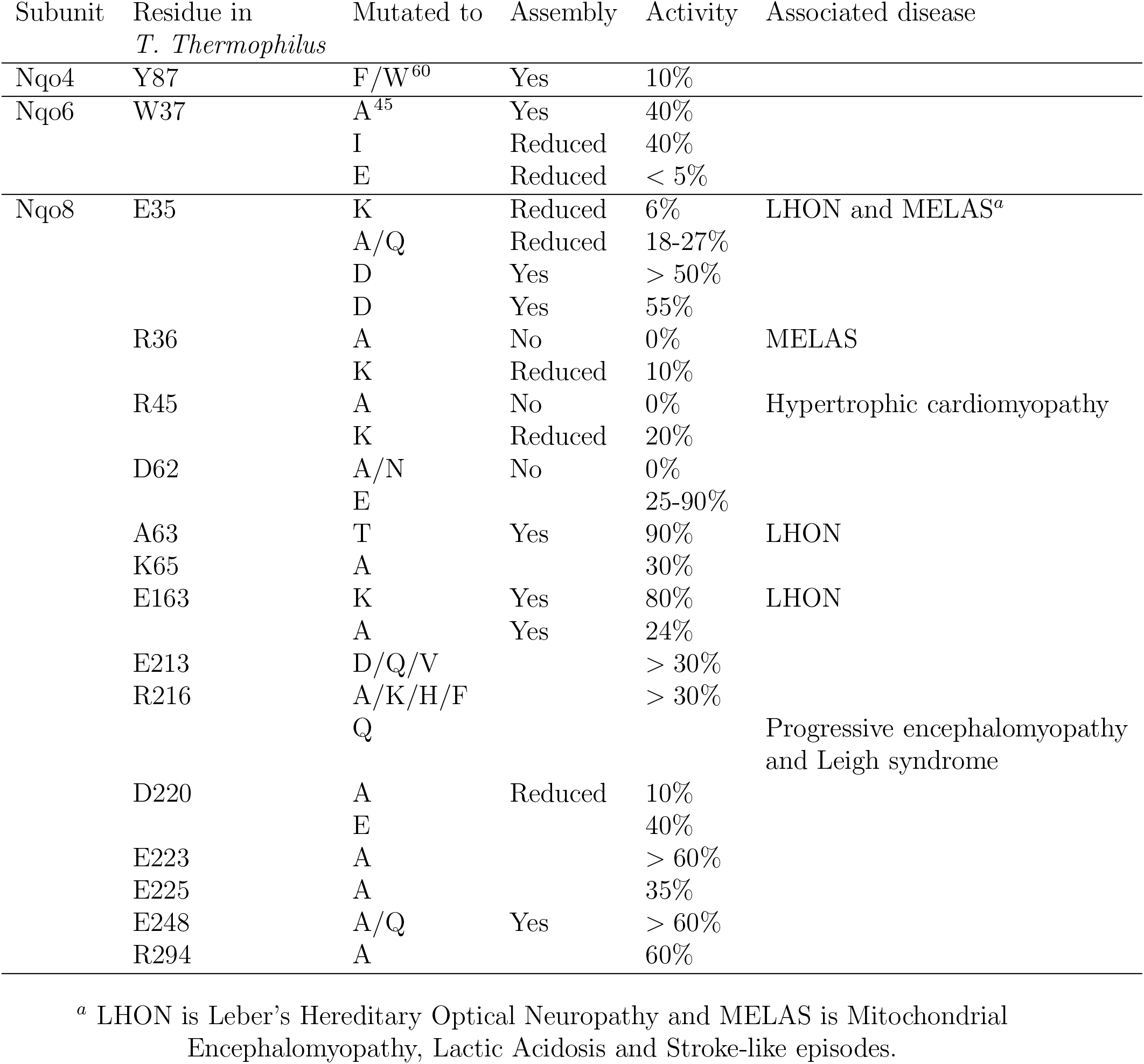
Point mutations in residues near the Q-chamber with the assembly of the mutated protein complex (or content of complex I), enzymatic activity of the mutant in relation to wild-type and associated human disease. Data taken from,^8^ except when explicitly referenced.

## References

(1) Brandt, U. Energy Converting NADH: Quinone Oxidoreductase (Complex I). Annu. Rev. Biochem. 2006, 75, 69–92.

(2) Hirst, J. Mitochondrial Complex I. Annu. Rev. Biochem. 2013, 82, 551–575.

(3) Verkhovskaya, M.; Bloch, D. A. Energy-converting respiratory Complex I: 0n the way to the molecular mechanism of the proton pump. Int. J. Biochem. Cell Biol. 2013, 45, 491–511.

(4) Sazanov, L. A. A giant molecular proton pump: structure and mechanism of respiratory complex I. Nature Rev. Mol. Cell Biol. 2015, 16, 375–388.

(5) Wikstrom, M.; Sharma, V.; Kaila, V. R. I.; Hosler, J. P.; Hummer, G. New Perspectives on Proton Pumping in Cellular Respiration. Chem. Rev. 2015. 115, 2196–2221.

(6) Pätsi, J.; Maliniemi, P.; Pakanen, S.; Hinttala, R.; Uusimaa, J.; Majamaa, K.; Nystrom, T.; Kervinen, M.; Hassinen, I. E. LHON/MELAS overlap mutation in ND1 subunit of mitochondrial complex I affects ubiquinone binding as revealed by modeling in *Escherichia coli* NDH-1. Biochim. Biophys. Acta 2012, 1817, 312–318.

(7) Cadenas, S. Mitochondrial uncoupling, ROS generation and cardioprotection. Biochim. Biophys. Acta 2018, 1859, 940–950.

(8) Baradaran, R.; Berrisford, J. M.; Minhas, G. S.; Sazanov, L. A. Crystal structure of the entire respiratory complex I. Nature 2013, 494, 443–448.

(9) Zickermann, V.; Wirth, C.; Nasiri, H.; Siegmund, K.; Schwalbe, H.; Hunte, C.; Brandt, U. Mechanistic insight from the crystal structure of mitochondrial complex I. Science 2015, 347, 44–49.

(10) Zhu, J.; Vinothkumar, K. R.; Hirst, J. Structure of mammalian respiratory complex I. Nature 2016, 536, 354–358.

(11) Fiedorczuk, K.; Letts, J. A.; Deglies-posti, G.; Kaszuba, K.; Skehel, M.; Sazanov, L. A. Atomic structure of the entire mammalian mitochondrial complex I. Nature 2016, 538, 406–410.

(12) Wu, M.; Gu, J.; Guo, R.; Huang, Y.; Yang, M. Structure of mammalian respiratory supercomplex *I_1_III_2_IV_1_*. Cell 2016, 167, 1598–1609.

(13) Agip, A.-N. A.; Blaza, J. N.; Bridges, H. R.; Viscomi, C.; Rawson, S.; Muench, S. P.; Hirst, J. Cryo-EM structures of complex I from mouse heart mitochondria in two biochemically defined states. Nat. Struct. Mol. Biol. 2018, 25, 548–556.

(14) Gumbart, J. C.; Teo, I.; Roux, B.; Schulten, K. Reconciling the roles of kinetic and thermodynamic factors in membrane-protein insertion. J. Am. Chem. Soc. 2013, 135, 2291–2297.

(15) Bai, F.; Xu, Y.; Chen, J.; Liu, Q.; Gu, J.; Wang, X.; Ma, J.; Li, H.; Onuchic, J. N.; Jiang, H. Free energy landscape for the binding process of Huperzine A to acetylcholinesterase. Proc. Natl. Acad. Sci. USA 2013, 110, 4273–4278.

(16) Bisha, I.; Rodriguez, A.; Laio, A.; Magistrato, A. Metadynamics simulations reveal a Na+ independent exiting path of galactose for the inward-facing conformation of vSGLT. PLoS Comput. Biol. 2014, 10, 1–8.

(17) Paloncyová, M.; V., N.; Berka, K.; Laio, A.; Otyepka, M. Role of enzyme flexibility in ligand access and egress to active site: bias-exchange metadynamics study of 1,3,7-trimethyluric acid in cytochrome P450 3A4. J. Chem. Theory Comput. 2016, 12, 2101–2109.

(18) Torabifard, H.; Cisneros, G. A. Computational investigation of O2 diffusion through an intra-molecular tunnel in AlkB. Influence of polarization on O_2_ transport. Chem. Sci. 2017, 8, 6230–6238.

(19) Nunes-Alves, A.; Zuckerman, D. M.; Arantes, G. M. Escape of a small molecule from inside T4 lysozyme by multiple pathways. Biophys. J. 2018, 114, 1058–1066.

(20) Van Eerden, F. J.; Melo, M. N.; Frederix, P. W.; Periole, X.; Marrink, S. J. Exchange pathways of plastoquinone and plastoquinol in the photosystem II complex. Nat. Commun. 2017, 8, 1–8.

(21) Warnau, J.; Sharma, V.; Gamiz-Hernandez, A. P.; Luca, A. D.; Haapanen, O.; Vattulainen, I.; Wikstrom, M.; Hummer, G.; Kaila, V. R. I. Redox-coupled quinone dynamics in the respiratory complex I. Proc. Natl. Acad. Sci. USA 2018, 115, E8413–E8420.

(22) Galassi, V. V.; Arantes, G. M. Partition, orientation and mobility of ubiquinones in a lipid bilayer. Biochim. Biophys. Acta 2015, 1847, 1345.

(23) Sali, A.; Blundell, T. L. Comparative protein modelling by satisfaction of spatial restraints. J. Mol. Biol. 1993, 234· 779–815.

(24) Sharma, V.; Belevich, G.; Gamiz-Hernandez, A. P.; Róg, T.; Vattulainen, I.; Verkhovskaya, M. L.; Wiksträm, M.; Hummer, G.; Kaila, V. R. I. Redox-induced activation of the proton pump in the respiratory complex I. Proc. Natl. Acad. Sci. USA 2015, 112, 11571–11576.

(25) Huey, R.; Morris, G. M.; Olson, A. J.; Goodsell, D. S. A Semiempirical Free Energy Force Field with Charge-Based Desolvation. J. Comput. Chem. 2007. 28, 1145–1152.

(26) Javanainen, M. Universal Method for Embedding Proteins into Complex Lipid Bilayers for Molecular Dynamics Simulations. J. Chem. Theory Comput. 2014, 10, 2577–2582.

(27) Klauda, J. B.; Venable, R. M.; Freites, J. A.; O‘Connor, J. W.; Tobias, D. J.; Mondragon-Ramirez, C.; Vorobyov, I.; Jr., A. D. M.; Pastor, R. W. Update of the CHARMM AllAtom Additive Force Field for Lipids: Validation on Six Lipid Types. J. Phys. Chem. B 2010, 114, 7830–7843.

(28) Huang, J.; MacKerell Jr, A. D. CHARMM36 all-atom additive protein force field: Validation based on comparison to NMR data. J. Comp. Chem. 2013, 34, 2135–2145.

(29) Jorgensen, W. L.; Chandrasekhar, J.; Madura, J. D.; Impey, R. W.; Klein, M. L. Comparison of simple potential functions for simulating liquid water. J. Chem. Phys. 1983, 79, 926–935.

(30) Chang, C. H.; Kim, K. Density Functional Theory Calculation of Bonding and Charge Parameters for Molecular Dynamics Studies on [FeFe] Hydrogenases. J. Chem. Theory Comput. 2009, 5, 1137–1145.

(31) McCullagh, M.; Voth, G. A. Unraveling the role of the protein environment for [FeFe]-hydrogenase: A new application of coarse-graining. J. Phys. Chem. B 2013, 117, 4062–4071.

(32) Abraham, M. J.; Murtola, T.; Schulz, R.; Pall, S.; Smith, J. C.; Hess, B.; Lindahl, E. GR0MACS: High performance molecular simulations through multi-level parallelism from laptops to supercomputers. Software X 2015, 1–2, 19–25.

(33) Darden, T.; York, D.; Pedersen, L. Particle mesh Ewald: An N·log(N) method for Ewald sums in large systems. J. Chem. Phys. 1993, 98, 10089–10092.

(34) The PyMOL Molecular Graphics System, Version 1.8; Schrodinger, LLC, 2015.

(35) Hunter, J. D. Matplotlib: A 2D graphics environment. Comput. Sci. Eng. 2007, 9, 90–95.

(36) Branduardi, D.; Gervasio, F. L.; Parrinello, M. From A to B in free energy space. J. Chem. Phys. 2007, 126, 054103.

(37) Tribello, G. A.; Bonomi, M.; Branduardi, D.; Camilloni, C.; Bussi, G. Plumed 2: New feathers for an old bird. Comput. Phys. Commun. 2014, 185, 604–613.

(38) Roux, B. The calculation of the potential of mean force using computer simulations. Comput. Phys. Commun. 1995, 91, 275–282.

(39) Field, M. J. The pDynamo Program for Molecular Simulations using Hybrid Quantum Chemical and Molecular Mechanical Potentials. J. Chem. Theory Comput. 2008, 4, 1151–1161.

(40) Johnson, R. W. An introduction to the bootstrap. Teaching Statistics 2001, 23, 49–54.

(41) Gamiz-Hernandez, A. P.; Jussupow, A.; Johansson, M. P.; Kaila, V. R. Terminal Electron-Proton Transfer Dynamics in the Quinone Reduction of Respiratory Complex I. J. Am. Chem. Soc. 2017, 139, 16282–16288.

(42) Fato, R.; Estornell, E.; Di Bernardo, S.; Pallotti, F.; Castelli, G. P.; Lenaz, G. Steady-state kinetics of the reduction of coenzyme Q analogs by complex I (NADH:ubiquinone oxidoreductase) in bovine heart mitochondria and submito-chondrial particles. Biochemistry 1996, 35, 2705–2716.

(43) Zickermann, V.; Barquera, B.; Wiksträm, M.; Finel, M. Analysis of the pathogenic human mitochondrial mutation ND1/3460, and mutations of strictly conserved residues in its vicinity, using the bacterium Paracoccus denitrificans. Biochemistry 1998, 37, 11792–11796.

(44) Sakamoto, K.; Miyoshi, H.; Ohshima, M.; Kuwabara, K.; Kano, K.; Akagi, T.; Mogi, T.; Iwamura, H. Role of the isoprenyl tail of ubiquinone in reaction with respiratory enzymes: studies with bovine heart mitochondrial complex I and *Escherichia coli* bo-type ubiquinol oxidase. Biochemistry 1998, 37, 15106–15113.

(45) Angerer, H.; Nasiri, H. R.; Niedergesaβ, V.; Kerscher, S.; Schwalbe, H.; Brandt, U. Tracing the tail of ubiquinone in mitochondrial complex I. Biochim. Biophys. Acta 2012, 1817, 1776–1784.

(46) Snyder, P. W.; Mecinovic, J.; Moustakas, D. T.; III, S. W. T.; Harder, M.; Mack, E. T.; Lockett, M. R.; Hroux, A.; Sherman, W.; Whitesides, G. M. Mechanism of the hydrophobic effect in the biomolecular recognition of arylsulfonamides by carbonic anhydrase. Proc. Natl. Acad. Sci. USA 2011, 108, 17889–17894.

(47) Yao, H.; Ke, H.; Zhang, X.; Pan, S.-J.; Li, M.-S.; Yang, L.-P.; Schrecken-bach, G.; Jiang, W. Molecular Recognition of Hydrophilic Molecules in Water by Combining the Hydrophobic Effect with Hydrogen Bonding. J. Am. Chem. Soc. 2018, 140, 13466–13477.

(48) Fedor, J. G.; Jones, A. J. Y.; Luca, A. D.; Kaila, V. R. I.; Hirst, J. Correlating kinetic and structural data on ubiquinone binding and reduction by respiratory complex I. Proc. Natl. Acad. Sci. USA 2017, 114, 12737–12742.

(49) Zhang, L.; Hermans, J. Hydrophilicity of cavities in proteins. Proteins: Struct. Funct. Genet. 1996, 24, 433–438.

(50) Persson, F.; Halle, B. Transient Access to the Protein Interior: Simulation versus NMR. J. Am. Chem. Soc. 2013, 135, 8735–8748.

(51) Di Luca, A.; Gamiz-Hernandez, A. P.; Kaila, V. R. I. Symmetry-related proton transfer pathways in respiratory complex I. Proc. Natl. Acad. Sci. USA 2017, 114, 6314–6321.

(52) Isom, D. G.; Castaneda, C. A.; Cannon, B. R.; Velu, P. D.; García-Moreno E., B. Charges in the hydrophobic interior of proteins. Proc. Natl. Acad. Sci. USA 2010, 107, 16096–16100.

(53) Moser, C. C.; Keske, J. M.; Warncke, K.; Farid, R. S.; Dutton, P. L. Nature of Biological Electron Transfer. Nature 1992, 355, 796–802.

(54) Ohnishi, T. Iron-sulfur clusters & semiquinones in Complex I. Biochim. Biophys. Acta 1998, 1364, 186–206.

(55) Ohnishi, T.; Sled, V. D.; Yano, T.; Yagi, T.; Burbaev, D. S.; Vinogradov, A. D. Structure-function studies of iron-sulfur clusters and semiquinones in the NADH-Q oxidoreductase segment of the respiratory chain. Biochim. Biophys. Acta 1998, 1365, 301–308.

(56) Yano, T.; Dunham, W. R.; Ohnishi, T. Characterization of the *Δμ_H_* +-sensitive ubisemiquinone species (SQ_Nf_) and the interaction with cluster N2: New insight into the energy-coupled electron transfer in complex I. Biochemistry 2005, 44, 1744–1754.

(57) Hirst, J.; Roessler, M. M. Energy conversion, redox catalysis and generation of reactive oxygen species by respiratory complex I. Biochim. Biophys. Acta 2016, 1857, 872–883.

(58) Kurki, S.; Zickermann, V.; Kervinen, M.; Hassinen, I.; Finel, M. Mutagenesis of three conserved Glu residues in a bacterial homologue of the ND1 subunit of complex I affects ubiquinone reduction kinetics but not inhibition by dicyclohexylcarbodiimide. Biochemistry 2000, 39, 13496–13502.

(59) Fersht, A. Structure and Mechanism in Protein Science: A Guide to Enzyme Catalysis and Protein Folding, 1st ed.; W. H. Freeman: New York, 1999.

(60) Tocilescu, M. A.; Fendel, U.; Zwicker, K.; Drose, S.; Kerscher, S.; Brandt, U. The role of a conserved tyrosine in the 49-kDa subunit of complex I for ubiquinone binding and reduction. Biochim. Biophys. Acta 2010, 1797, 625–632.

(61) Pomorski, T. G.; Menon, A. K. Lipid somersaults: Uncovering the mechanisms of protein-mediated lipid flipping. Prog. Lipid Res. 2016, 64, 69–84.

(62) Furuhashi, M.; Hotamisligil, G. S. Fatty acid-binding proteins: role in metabolic diseases and potential as drug targets. Nat. Rev. Drug Discov. 2008, 7, 489–503.

(63) Bussi, G.; Donadio, D.; Parrinello, M. Canonical sampling through velocity rescaling. J. Chem. Phys. 2007, 126, 014101.

(64) Berendsen, H. J. C.; Postma, J. P. M.; van Gunsteren, W. F.; DiNola, A.; Haak, J. R. Molecular dynamics with coupling to an external bath. J. Chem. Phys. 1984, 81, 3684–3690.

(65) Hess, B.; Bekker, H.; Berendsen, H. J. C.; Fraaije, J. G. E. M. LINCS: A linear constraint solver for molecular simulations. J. Comput. Chem. 1997, 18, 1463–1472.

(66) Woo, H.-J.; Roux, B. Calculation of absolute protein-ligand binding free energy from computer simulations. Proc. Natl. Acad. Sci. USA 2005, 102, 6825–6830.

(67) Blaza, J. N.; Vinothkumar, K. R.; Hirst, J. Structure of the Deactive State of Mammalian Respiratory Complex I. Structure 2018, 26, 312–319.

(68) Ashkenazy, H.; Abadi, S.; Martz, E.; Chay, O.; Mayrose, I.; Pupko, T.; BenTal, N. ConSurf 2016: an improved methodology to estimate and visualize evolutionary conservation in macromolecules. Nucl. Acids Res. 2016, 53, 199–206.

